# Restoring Marčenko-Pastur noise estimation and denoising of diffusion MRI under common image reconstruction

**DOI:** 10.64898/2026.11.12.766195

**Authors:** Laurin Mordhorst, Karsten Tabelow, Siawoosh Mohammadi

## Abstract

Diffusion magnetic resonance imaging can reveal microstructure in the living brain, but the measurements are inherently noisy. Marčenko–Pastur denoising is a widely used remedy, helping to separate signal from noise under a simplified noise covariance model. However, we find that the model breaks under common image reconstructions across 28 public in vivo datasets, resulting in weaker denoising than intended. We restore the intended denoising behavior with a noise covariance model estimated from the reconstructed images, and validate the estimated model on repeated ex vivo and in vivo scans. The method requires no additional measurements or knowledge of the reconstruction, opening headroom for microstructure analyses in both existing and newly acquired data.

## 1 Introduction

Brain tissue has a complex microstructure, whose composition and organization vary across regions (Amunts & Zilles, 2015; Axer et al., 2011; Mordhorst et al., 2026; Ruthig et al., 2024; Shapson-Coe et al., 2024). Diffusion-weighted magnetic resonance imaging (dMRI) can probe tissue microstructure in the living brain non-invasively (Basser et al., 1994; Jensen et al., 2005; Mordhorst et al., 2025; Palombo et al., 2020; Uhl et al., 2024). Its sensitivity to microstructure is often limited by noise: diffusion weighting produces contrast by attenuating the signal, yielding a low signal-to-noise ratio (SNR). The SNR is set by the acquisition protocol, where it can only be traded for scan time or resolution. Denoising sits largely outside this trade, raising the SNR after the scan, and has therefore become a standard processing step (David et al., 2024; Garyfallidis et al., 2014; Manzano Patron et al., 2024; Schilling et al., 2026; Tax et al., 2022; Tournier et al., 2019).

Denoising methods exploit the redundancy of dMRI acquisitions, which sample the same underlying signal profile across different diffusion weightings. One line of work can stay within a single voxel, predicting the voxel’s signal at each diffusion weighting from the signal at the remaining diffusion weightings (Fadnavis et al., 2020). Others additionally assume similar signal profiles across different voxels (Becker et al., 2014; Coupe et al., 2008; Manjón et al., 2010), so that a few patterns describe a whole group of them, usually a local patch. Predominantly, these patterns are estimated via principal component analysis (PCA), and the remaining components are discarded as noise (Manjón et al., 2013; Manzano Patron et al., 2024).

The difficulty in PCA denoising is determining the threshold that separates signal from noise. Marčenko–Pastur (MP) denoising takes it from random matrix theory. The component variances form a spectrum that follows the MP distribution (Marčenko & Pastur, 1967), whose sharp upper edge defines the threshold at a given noise level. That holds only if the noise is independent and identically distributed (i.i.d.) across the voxels of a patch. Thermal noise would satisfy this (Pruessmann et al., 1999; Robson et al., 2008), but denoising is typically applied after reconstruction, which reshapes the noise in two ways. First, the noise level can vary across voxels, breaking identical distribution, for example under parallel imaging (Breuer et al., 2009; Dietrich et al., 2007; Robson et al., 2008). Second, neighboring voxels can be correlated, breaking independence, for example after partial-Fourier reconstruction (Henriques et al., 2023; Noll et al., 1991). Both are described together by the noise covariance, which deforms the spectrum so the MP distribution no longer applies. Assuming it anyway misplaces the threshold, leading to weaker denoising than intended (Henriques et al., 2023; Moeller et al., 2021).

MP denoising approaches that require no additional data are most widely used (Manzano Patron et al., 2024; Schilling et al., 2026). The cost is that these methods need to estimate the noise level from the deformed spectrum (“MPPCA”, Veraart et al., 2016c; one variant of “NORDIC”, Moeller, 2026; Moeller et al., 2021), potentially misplacing the threshold even further than an exact noise level would. Additional measurements break this dependence, giving the noise level from repeated *b* = 0 volumes (Henriques et al., 2023) or a dedicated noise scan (Cordero-Grande et al., 2019; Moeller et al., 2021). In addition, NORDIC normalizes the spatially varying noise level for parallel imaging acquisitions, restoring part of the i.i.d. assumption. Cordero-Grande et al. (2019) model the full noise covariance, but require detailed knowledge of the reconstruction, which may not be possible for vendor reconstructions (D’Antonio et al., 2026).

Here, we propose a method to restore the i.i.d. noise assumption underlying MP denoising, without additional measurements or knowledge of the reconstruction. The core of our method is a noise covariance model with two components: a spatially varying noise level and a correlation pattern over the extent of a denoising patch. We assume the correlation to be stationary within the brain, because it arises from *k*-space operations such as partial Fourier, apodization and regridding, which affect every voxel alike (Cordero-Grande et al., 2019; Henriques et al., 2023; Veraart et al., 2016a). We estimate the model from the diffusion series itself, in an iterative loop where MP denoising and the noise covariance refine one another. We first illustrate in simplified simulations how noise covariance affects denoising. We then validate the proposed method in an ex vivo mouse brain, and demonstrate improved consistency of the denoised signal and of diffusion kurtosis imaging (DKI, Jensen et al., 2005) metrics across repeated scans and across scanners in vivo. Finally, we characterize the noise covariance across 28 public in vivo datasets spanning various scanners, acquisitions and reconstructions, suggesting that spatial correlation rather than a varying noise level affects patch-based MP denoising.

## 2 Methods

### 2.1 Marčenko–Pastur denoising under i.i.d. noise

#### Denoising

We consider patch-based denoising (Cordero-Grande et al., 2019; Henriques et al., 2023; Manjón et al., 2013; Moeller et al., 2021; Veraart et al., 2016c), where a sliding window sweeps the dMRI series voxel by voxel. Each voxel is denoised from a patch of *M* neighboring voxels around it, typically an isotropic cuboid with 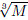 voxels per side. Let *X* ∈ ℝ*^M^*^×^*^N^* hold one patch, with its *M* voxels along the rows and the *N* dMRI volumes of the acquisition along the columns. We write all expressions below for *M* ≥ *N*. We model *X* = *S* + *E*, with an approximately low-rank signal *S* ∈ ℝ*^M^*^×^*^N^* of rank *P* ≪ min(*M, N*) and additive noise *E* ∈ ℝ*^M^*^×^*^N^*, whose columns share the patch noise covariance Σ ∈ R*^M^*^×^*^M^*. Denoising is then a projection onto the span of the *P* leading right singular vectors of *X*, collected in *V_P_*∈ ℝ*^N^*^×^*^P^*, equivalently the leading eigenvectors of *X*^⊤^*X*,

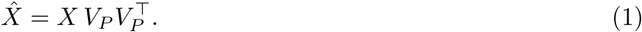

The difficulty is determining *P*, which random matrix theory helps resolve. If the entries of *E* are independent and identically distributed with variance *σ*^2^ across volumes and across the voxels of a patch, so that Σ = *σ*^2^*I* with the identity *I* ∈ ℝ*^M^*^×^*^M^*, the noise produces an eigenvalue bulk that follows the MP distribution (Marčenko & Pastur, 1967). Components above the bulk’s approximate upper edge 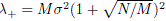 are then taken as signal.

#### Estimating the noise level σ

In practice, *σ* is typically unknown. MPPCA (Veraart et al., 2016c) estimates *σ* from the same eigenvalue spectrum it is thresholding, together with *P*. For a candidate rank *p*, the *N p* smallest eigenvalues *λ_p_*_+1_, …, *λ_N_* of *X*^⊤^*X*, ordered as *λ*_1_≥ … ≥ *λ_N_*, are taken as noise, and their suitably normalized mean and spread each provide an estimate of *σ*^2^ (Cordero-Grande et al., 2019). At the true rank *p* = *P*, these eigenvalues should follow the MP distribution, so the two estimates agree approximately. MPPCA therefore increases *p* from zero until the spread estimate falls below the mean estimate, and determines the noise level from the mean at that rank,

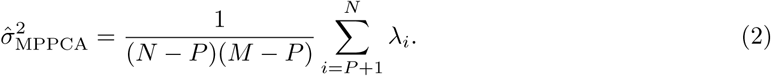

Despite assuming i.i.d. noise within each patch, MPPCA yields a spatial noise map *σ*^_MPPCA_(*x*), with *x* a voxel position, which captures variation coarser than a patch (Veraart et al., 2016b).

### 2.2 Marčenko–Pastur denoising under non-i.i.d. noise

Reconstruction breaks the i.i.d. noise assumption of MP denoising, varying the noise level across voxels (Breuer et al., 2009; Dietrich et al., 2007; Robson et al., 2008) and correlating neighboring ones (Henriques et al., 2023; Noll et al., 1991). The patch noise covariance Σ is then unlike *σ*^2^*I*. With known Σ, the i.i.d. noise assumption can be restored by whitening each patch prior to MP denoising. While Σ is typically unknown, we propose that it can be estimated from the components MPPCA discards as noise in each patch in an iterative approach (see Figure 1). A better Σ lets MPPCA split signal from noise more accurately, and a better split gives a better Σ. Hence, we iterate to the fixed point, where Σ predicts the covariance of the components MPPCA discards as noise under it. The approach is detailed below.

**Figure 1:**
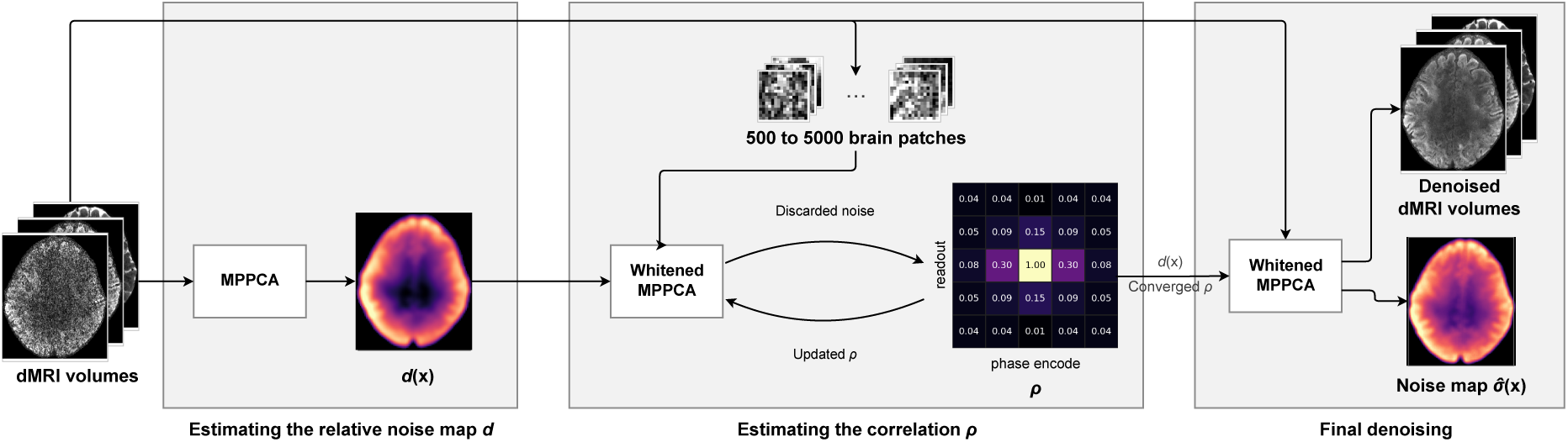
Marčenko–Pastur denoising with a noise covariance model. To estimate the relative noise map *d*(*x*), MPPCA (Veraart et al., 2016c) runs on the dMRI volumes. Then, the correlation *ρ* is estimated from a whole brain subset of 500 to 5000 patches. To this end, we alternate whitened MPPCA, which transforms each patch with the whitening matrix *W* built from *d*(*x*) and *ρ* before a regular MPPCA call (see Eq. (3)), with a refit of *ρ* to the components MPPCA discards as noise. The panel illustrates *ρ* along readout and phase encode. After convergence of *ρ*, we denoise the image with one whitened MPPCA pass, giving the denoised dMRI volumes and the noise level map *σ*^(*x*) of Eq. (6).

#### Pre-whitening

The i.i.d. noise assumption can be restored by transforming a patch with a suitable whitening matrix *W* (Gavish et al., 2023; Leeb & Romanov, 2021), so that the patch noise covariance becomes *W* Σ*W* ^⊤^ = *σ*^2^*I*. In this convention, *σ* stays in the patch for MPPCA to estimate. The truncation of Eq. (1) then becomes

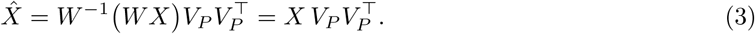

Here *W* cancels from the output, but it changes the retained subspace, i.e., *V_P_* now holds the leading right singular vectors of *WX* rather than of *X*.

#### Noise covariance model

We model the patch noise covariance Σ ∈ ℝ*^M^*^×*M*^ as a spatially varying noise level, split into the scalar *σ* and its relative variation in a diagonal matrix *D* ∈ ℝ*^M^*^×*M*^, and a correlation matrix *C* ∈ ℝ*^M^*^×*M*^:

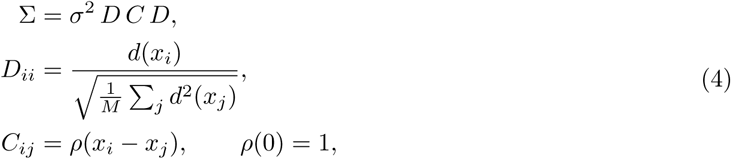

with *x_i_* and *x_j_* the positions of the patch’s voxels *i* and *j*. *D* holds the patch’s values of a whole-image relative noise map *d*, normalized to unit root mean square over the patch, so that *σ*^2^ is the mean noise variance over the patch. We assume the correlation to be stationary and shared across patches, so that the lag function *ρ* gives it from the displacement alone. The stationarity assumption rests on the correlation arising largely from partial Fourier, apodization and gridding, which act in *k*-space and therefore induce nearly shift-invariant correlations (Cordero-Grande et al., 2019; Henriques et al., 2023; Veraart et al., 2016a). We assume this within the brain, where the remaining steps of the reconstruction leave the noise additive. The parameterization of *C* by *ρ* greatly reduces the number of free parameters: for example, at a typical patch size of 5×5×5, from 7750 correlations per patch to 364 shared across the whole brain.

#### Estimating the relative noise map d

We estimate *d* once, before the iterative approach that fits *ρ*, from the noise level MPPCA estimates without whitening. To this end, we run MPPCA on patches centered at every second voxel along each axis, assign each patch’s noise variance to all of its voxels, average where patches overlap and take the square root to convert the noise variance into a noise level. The averaging smooths the variability that MPPCA’s per-patch rank selection leaves in the noise level map, trading spatial detail for a stable *d*. In contrast to a NORDIC variant, which also estimates a noise level map with MPPCA before denoising (Moeller, 2026; Moeller et al., 2021), we only need *d* to approximate the relative spatial variation of *σ*(*x*), not its absolute scale.

#### Estimating the correlation ρ

Keeping *d* fixed, we estimate *ρ* by alternating with MPPCA. MPPCA needs *ρ* to whiten each patch before splitting it into signal and noise, and *ρ* is fitted to the noise that split discards (see Figure 1). We initialize *ρ* as white noise, unit at zero displacement and zero elsewhere. Each iteration starts by whitening every patch with *W* = *C*^−1*/*2^*D*^−1^, built from the current *ρ* and the known *d*, and running MPPCA on the result, yielding the retained rank *P* and the leading *P* left singular vectors *U_P_* of *WX*. The discarded noise contains information about *C*, but its covariance is not *C* itself, because MPPCA also removes noise along the retained signal components. We therefore fit *C* to the discarded noise under the projection *Q* = *I U_P_ U* ^⊤^ that produced it. Holding the split fixed, we approximate the *N P* nonzero components of the discarded noise *QWX* as independent and Gaussian with covariance *v QWDĈDW* ^⊤^*Q*^⊤^, where *v* is the patch’s noise variance and *Ĉ* the correlation sought. The noise variance differs between patches. For a given *Ĉ*, we therefore replace each *v* by the value that maximizes the likelihood, which has a closed form, leaving a log-likelihood in *Ĉ* alone. Assuming a single *Ĉ* across patches, we sum the log-likelihood over patches and restrict *Ĉ* to the stationary form of Eq. (4), one unknown per displacement, giving

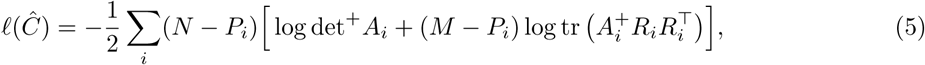

with *i* indexing patches, *R_i_* = *Q_i_WX_i_* the discarded noise and *A_i_* = *Q_i_WDĈDW* ^⊤^*Q_i_*^⊤^ its covariance up to the patch noise variance. The projection makes *A_i_* singular, so det^+^ and *A_i_*^+^ are the determinant and the inverse on the *M – P_i_* dimensional range of *Q_i_*, where *A_i_* is invertible. We maximize the log-likelihood by Fisher scoring (Nocedal & Wright, 2006), taking one step per iteration. A backtracking line search halves the step until the log-likelihood rises by a fixed share of the predicted amount, the Armijo condition, and *Ĉ* remains a valid correlation. One direction is barely visible to the fit, a blend of *Ĉ* toward the all-ones matrix **11**^⊤^, with **1** ∈ ℝ*^M^* the all-ones vector, which the profiled patch variance absorbs and which adds covariance along *D***1**. For a flat *d* this is the flat mode *u* ∈ ℝ*^M^* with 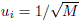. MPPCA generally retains the flat mode as signal, since a patch of image intensity is dominated by its mean, so the discarded noise constrains this direction only weakly. We fix the direction by requiring *ρ* to average to zero over the largest displacements the patch spans, assuming the correlation decays with distance. The fit uses 500 to 5000 patches sampled across the brain, non-overlapping where the brain includes enough voxels. The number of patches is generous, as few patches would suffice to estimate the correlation, whereas the number of volumes has more influence on the estimated correlation (see Figure S1). We stop once a full step moves no value of *ρ* by more than 5 × 10^−4^, and after at most 20 iterations.

#### Denoising

With all parameters estimated, we apply a final pass over all patches of the image. We whiten each patch with *W* = *C*^−1*/*2^*D*^−1^ and truncate as in Eq. (3). MPPCA fits the threshold per patch on the whitened patch, with nothing supplied to it, so the method reduces to it exactly when *C* = *I* and *d* is flat.

#### Estimating the noise level σ

The final pass also yields *σ*^_MPPCA_ per patch, giving the patch noise covariance 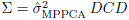. While *σ*^_MPPCA_ could serve as the reported noise level, we instead derive it from the estimated covariance, excluding the flat mode *u*. We report an effective noise variance for the center voxel, excluding the flat mode,

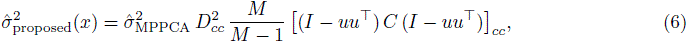

with *c* the index of the patch’s center voxel, at position *x*, so that *D* and the projected correlation are both read at that voxel’s diagonal entry. This equals *σ*^2^ for i.i.d. noise. Under predominantly positive spatial correlation, it falls below the voxel’s own noise variance Σ*_cc_*. This choice affects only the reported noise level, not the denoised images, which depend on *W* and the retained rank alone.

### 2.3 Simulations

We studied how a noise covariance affects MP denoising in simplified dMRI simulations on a subject phantom.

#### Subject phantom

We built a subject phantom from the 15 central slices of the complex average of six repeats of a 2 mm isotropic UK Biobank protocol (Dataset A of Manzano Patron et al., 2024, 2026). The data come preprocessed and averaged, with the repeats rotated to the real axis by their estimated phase (Moeller et al., 2021) and corrected for motion, susceptibility and eddy current distortions (Andersson & Sotiropoulos, 2016; Andersson et al., 2003; Mohammadi-Nejad et al., 2023). We fitted the signal of every voxel onto a spherical-harmonic basis of even degree up to four, per diffusion-weighting *b*. We then generated the phantom by evaluating the fit at the protocol’s gradient directions. Averaging over repeats and the low-order spherical-harmonic fit both cost spatial detail, which we accepted for a low residual noise covariance and for meeting the MP low-rank assumption, allowing us to study the effect of an imposed noise covariance in isolation.

#### Simulating reconstruction effects

We simulated effects standing in for the *k*-space weighting (Chen et al., 2026; Tax et al., 2022; Veraart et al., 2016a) and the coil combination (Moeller et al., 2021) of a real reconstruction, simplified so that each factor can be swept on its own. To this end, we generated noisy signals 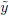 from the noise-free *y* = *ℱ*^−1^ {*w ℱs}* and white Gaussian noise *ε* of unit variance, drawn per voxel and per volume, as *ŷ* = *y* + *σ*_thermal_ *d ℱ*^−1^{*w ℱε}*. Here, *s* ∈ ℝ*^nx^*^×^*^ny^* ^×^*^nz^* ^×^*^N^* is the subject phantom over *n_x ×_n_y_* in-plane voxels, *n_z_* slices and *N* diffusion volumes, *ℱ*the in-plane discrete Fourier transform, *w* ∈ ℝ*^nx^*^×^*^ny^* a *k*-space window imposing correlation between voxels, *d* ∈ ℝ*^nx^*^×^*^ny^* ^×^*^nz^* the spatially varying relative noise level, and *σ*_thermal_ ∈ ℝ the thermal noise level entering the reconstruction. We realized *w* as separable in plane, applying the same blend between a flat and a Hann window, (1 – *f_C_*) + *f_C_* cos^2^(*πk/n*), along each axis, where *f_C_* sets the correlation strength, *k* the *k*-space sample index measured from the center and *n* the number of samples along that axis. For *d*, we took a fixed smooth field that varies over a few voxels (see Figure 2a) and rescaled it over the brain to [1, 1 + *f_d_*], so that *f_d_* sets the maximum noise amplification, as under parallel imaging (Breuer et al., 2009; Dietrich et al., 2007; Robson et al., 2008). The ground truth image noise level is then *σ*(*x*) = *σ*_thermal_ *d*(*x*) rms(*w*), where the root mean square (rms) of the window over the *k*-space samples measures how much noise survives the filtering.

**Figure 2:**
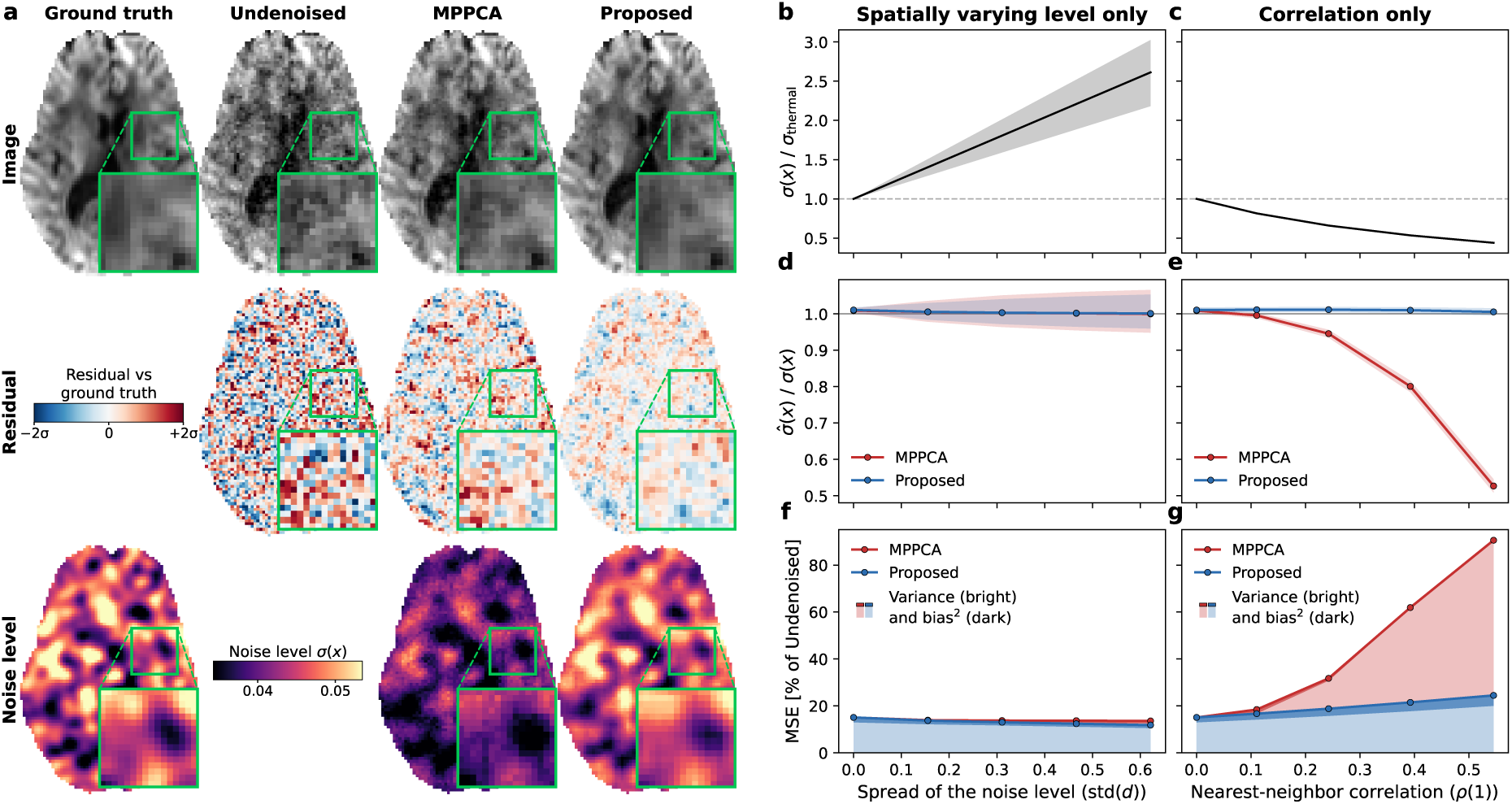
Simulations of reconstruction effects and their impact on denoising. **(a)** Qualitative comparison of denoising methods against the ground truth. Rows show a diffusion-weighted slice at *b* = 2000 s*/*mm^2^, the residual against the ground truth, and the post-reconstruction noise level *σ*(*x*). Insets magnify the boxed region. **(b–g)** Quantitative metrics for simulations under noise with a spatially varying level alone (b, d, f) and with a spatial correlation alone (c, e, g), swept by the spatial spread of the noise level std(*d*) and by the correlation of a voxel with its nearest neighbor *ρ*(1) (see Eq. (4)). **(b–c)** Post-reconstruction noise level *σ*(*x*) relative to the constant thermal noise level *σ*_thermal_, as the median across voxels with bands indicating the interquartile range. **(d–e)** Each denoising method’s noise level estimate *σ*^(*x*) over *σ*(*x*), as the median across voxels and noise draws, with bands indicating the interquartile range. **(f–g)** Mean squared error (MSE) of the denoised series against the ground truth, relative to the undenoised series’ MSE, with shading splitting it into the variance across noise draws and the squared bias as the remainder. All data were generated at thermal SNR = 20. The data in (a) are one noise draw at *ρ*(1) = 0.32 and std(*d*) = 0.21 (*f_C_* = 0.5, *f_d_* = 1.0). The data in (b–g) are averaged over 20 noise draws per setting.

#### Evaluation

We used 20 noise draws per sweep setting at a thermal SNR of 20, with *f_C_* from 0 to 0.8 and *f_d_* from 0 to 3. We scored the estimated noise levels *σ*^_MPPCA_ and *σ*^_proposed_ against *σ*, as the median over voxels and noise draws, and each denoised series *ŷ* against the noise-free *y* over the brain voxels *i* of all volumes, by the mean squared error (MSE) ⟨(*ŷ_i_ y_i_*)^2^⟩*_i_*, normalized to the undenoised series’ error. We split the MSE into the variance across noise draws and the squared bias as the remainder.

### 2.4 Ex vivo mouse brain dMRI

We validated the method in an ex vivo mouse brain, where the absence of subject motion gives a reference for signal and noise.

#### Data

We used the publicly available ex vivo mouse brain magnitude dMRI data (Henriques et al., 2023), acquired on a preclinical 16.4 T Bruker Aeon Ascend system. The scan protocol used 30 gradient directions at each of *b* = 1000, 2000 and 3000 s*/*mm^2^, and 20 *b* = 0 images. The diffusion gradients lasted *δ* = 1.5 ms at a separation of Δ = 15 ms, at a TR of 3000 ms and a TE of 50 ms. The field of view was 12 mm × 12 mm on a 80 × 80 matrix, giving 150 µm × 150 µm in plane, over nine coronal slices of 0.7 mm. The EPI readout sampled during the gradient ramps and used a phase partial-Fourier factor of 6/8, reconstructed by zero-filling, so the noise is correlated. The sample was scanned at two averages and at twenty averages, the latter providing a high-SNR reference. The published volumes come corrected for spatial drift by a sub-pixel registration (Guizar-Sicairos et al., 2008).

#### Processing

We denoised the two-average dMRI volumes and fitted DTI and DKI (Basser et al., 1994; Garyfallidis et al., 2014; Henriques et al., 2021; Jensen et al., 2005). To obtain a high-SNR reference, we also fitted DTI and DKI on the undenoised twenty-average volumes. As a reference for the noise level, we took the standard deviation per voxel over the 20 undenoised *b* = 0 images of the two-average volumes. Since the standard deviation over 20 images is itself noisy, we pooled it over a 3^3^ neighborhood by the root mean square.

#### Evaluation

We scored the estimated noise levels *σ*^_MPPCA_ and *σ*^_proposed_ against the reference *σ* per brain voxel. At signal level we scored each dMRI series *ŷ* against the reference *y* over the brain voxels *i* of one shell, by the mean squared error 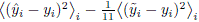, normalized to the undenoised series’ *ŷ* error. The subtracted term is the reference’s own noise variance, a tenth of the two-average series’ noise variance and therefore an eleventh of their squared difference. For the diffusion metrics, we scored each metric 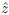 against the reference’s *z*, by the root mean square error relative to the reference metric’s mean, 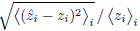 over white matter voxels *i* with fractional anisotropy above 0.25 in the twenty-average reference.

### 2.5 In vivo consistency within and across scanners

To validate the method in the human brain in vivo, we assessed repeatability, the agreement of the measurements across repeats on one scanner, and reproducibility, the agreement of the measurements across scanners in the same subjects.

#### Data

We used a subset of the ON-Harmony dataset (Warrington et al., 2023, 2025b), including magnitude dMRI from seven subjects and seven scanners of three vendors. Each subject has six repeats in separate sessions on one of the scanners, enabling a within-scanner comparison. For across-scanner comparison, each subject was scanned once on the other scanners of their cohort, where the two cohorts covered five and six of the seven scanners. All sessions of a subject lie days to months apart, at most about two years. The acquisition follows the UK Biobank diffusion protocol, with 50 directions each at *b* = 1000 s*/*mm^2^ and *b* = 2000 s*/*mm^2^ at 2 mm isotropic resolution. Resolution and *b*-shells are matched across the scanners, while partial Fourier, in-plane acceleration and multiband factor differ (see Table A1). Additionally, we used a four-repeat magnitude dMRI dataset at 0.9 mm isotropic resolution from one subject on a Siemens Prisma 3 T system (Dataset C of Manzano Patron et al., 2024, 2026). The data provide 199 volumes per repeat, 14 at *b* = 0, 93 at *b* = 1000 s*/*mm^2^ and 92 at *b* = 2000 s*/*mm^2^, acquired with a multiband factor of three and an in-plane acceleration factor of two. The four repeats were acquired back to back within one session.

#### Processing

We denoised the dMRI volumes with MPPCA and the proposed method. Then, we registered each subject’s repeats to a common template, correcting head position and geometric distortions, one template for the within-scanner and one for the across-scanner comparison. We built each as a multi-modal template, from the repeats’ DKI *S*_0_ and fractional anisotropy maps (Avants et al., 2008). We composed a single transform accounting for motion, susceptibility and eddy current distortions (Andersson & Sotiropoulos, 2016; Andersson et al., 2003; Jenkinson et al., 2012), and the registration to the template. We estimated this transform on the volumes denoised with MPPCA, in a forward pass that resamples at each stage. Then, we applied it in a single resampling step to undenoised and denoised volumes, and rotated the diffusion gradient directions accordingly. This approach minimizes interpolation and ensures that the same voxels are compared across denoising methods. We chose MPPCA to estimate the transform, profiting from improved estimation through denoising (Cieslak et al., 2024; Schilling et al., 2026) without favoring the proposed method. In the template space, we fitted DTI and DKI per repeat (Basser et al., 1994; Garyfallidis et al., 2014; Henriques et al., 2021; Jensen et al., 2005). We defined white matter from the tissue segmentation of a T1-weighted image (Avants et al., 2011; Lorio et al., 2016), registered to the diffusion template (Avants et al., 2008).

#### Evaluation

Within scanners, we scored each series by the unsupervised mean squared error (Morales et al., 2023) against the mean of the other repeats’ undenoised volumes, as MSE = ⟨(*a_ir_* – ⟨raw*_is_*⟩*_s≠r_*)^2^⟩ *_i,r_* – ⟨Var*_r_*(raw*_ir_*)⟩*_i_/*(*R –* 1), where *a_ir_* is the signal at voxel *i* in repeat *r*, raw*_ir_* the undenoised data and *R* the number of repeats. The second term accounts for the noise in the undenoised volumes. We normalized the MSE to the undenoised series’ error and split it into the variance across repeats, ⟨Var*_r_*(*a_ir_*)⟩*_i_*, and the squared bias as the remainder. Beforehand, we scaled each repeat by its mean *b* = 0 signal, so that overall intensity differences between repeats do not enter. For <u>diffusion</u> metrics, we scored each map *z* by its coefficient of variation (Ades-Aron et al., 2025), 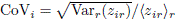, across the repeats within scanners and across the scanners, over white matter voxels *i*, where the T1-weighted segmentation gave a white matter posterior above 0.5.

### 2.6 The noise covariance across public datasets

To characterize the noise covariance and its effect on MP denoising across scanners, acquisitions and reconstructions, we evaluated 28 public human in vivo datasets.

#### Data

We screened OpenNeuro.org and the Human Connectome Project for datasets in the DTI and DKI regime, providing unprocessed post-reconstruction magnitude dMRI with at least 50 diffusion-weighted images at *b* ≤ 2500 s*/*mm^2^ and two subjects each. The datasets span several factors with potential impact on the noise covariance, including 12 scanner models from three vendors, partial-Fourier fractions from 0.75 to 1, in-plane acceleration up to 3, and multiband factors up to six. Table A2 lists dataset information.

#### Evaluation

We denoised all datasets and quantified the correction the proposed method applies to MPPCA’s noise level, *σ*^_proposed_*/σ*^_MPPCA_ 1. To attribute the correction to the components of the noise covariance model, we estimated the noise level (see Eq. (6)) again with the correlation alone and with the spatially varying noise level alone. We then expressed the resulting corrections as a share of the one the full model applies. In the absence of a reference, we scored the noise covariance Σ by the noise level it predicts, *σ*^_predicted_, for MPPCA’s reading, *σ*^_MPPCA_. To this end, we synthesized patches from the signal the proposed method keeps, with noise Σ^1*/*2^*ε*, where *ε* is white Gaussian of unit variance at the patch’s shape, and compared MPPCA’s median *σ*^ over 200 draws with the one on the real data. We pooled both over the patches by their root mean square and reported *σ*^_predicted_*/σ*^_MPPCA_ – 1. Further, we studied whether the correlation transfers across subjects. To this end, we synthesized noise with one subject’s correlation and whitened it with the second subject’s of the same dataset. Then, we evaluated whether the proposed method recovers the noise level it was synthesized with, *σ*^_proposed_*/σ –* 1. As a control for how specific the correlation is, we repeated the analysis with the mean correlation pattern over the panel, leaving out the dataset under evaluation.

## 3 Results

Across simulations and experimental data, we compare the proposed method against conventional MP-PCA, which runs without prior whitening. For both methods, we used the widely used patch size rule, the smallest odd isotropic patch with at least as many voxels as volumes (*M* ≥ *N*, Tournier et al., 2019). This benchmark allows us to assess the key contribution of the proposed method, the noise covariance model, in isolation.

### 3.1 Simulations

We simulated the impact of the noise covariance on denoising in a simplified setting, in which the noise covariance follows our proposed model and the MP low-rank assumption is well satisfied.

Figure 2a provides a qualitative comparison of a diffusion-weighted image, showing that both denoising methods improve on the undenoised data. Yet, MPPCA under-denoises and stays further away from the ground truth, leaving behind structured noise easily confused with anatomy rather than the per-voxel grain that residual i.i.d. noise would leave. Figure S2 further illustrates how much such structured noise varies between independent noise draws. The proposed method also reduces the residuals against the ground truth and better matches its noise level (see Figure 2a, center and bottom row). MPPCA underestimates the noise level, which translates into truncating too little noise, as observed qualitatively in the diffusion-weighted images.

Figure 2b–g assess the impact of the noise covariance quantitatively, sweeping its components in isolation. The simulated spatial noise variation raises the noise above the constant thermal noise level (see Figure 2b). In contrast, the correlation lowers it through the *k*-space weighting, which removes noise power and blurs the signal alike (see Figure 2c). The proposed method recovers the noise level accurately and keeps the mean squared error (MSE) of the signal low under both spatially varying noise and correlation, whereas MPPCA degrades steeply with increasing correlation (see Figure 2d–g). The gains of the proposed method arise largely from variance reduction, at slightly increased bias. The bias is similar across both methods under the setting MPPCA assumes, that is, the absence of correlation. Under correlation, the proposed method preserves a relatively stable bias, while MPPCA’s bias vanishes as it removes less and less of the noise. This is consistent with MPPCA falling short of its intended denoising under correlated noise, and consistent with the proposed method restoring it. While these results illustrate the impact of the noise covariance at a fixed number of volumes and patch size, their ratio further affects MPPCA’s performance (see Figure S3).

### 3.2 Ex vivo mouse brain dMRI

We validated the proposed method in an ex vivo mouse brain (Henriques et al., 2023), where twenty averages of the diffusion series give a reference for the signal and twenty consecutive *b* = 0 repeats give a reference for the noise level.

Figure 3a provides a qualitative comparison between diffusion-weighted images. While undenoised and denoised images hardly differ (top row), the difference maps against the reference make the improvement from denoising visible, with the proposed method yielding the smallest differences (center row). Notably, the noise level maps show the proposed method matching the overall scale of the reference much better than MPPCA (bottom row). Yet, neither method reproduces the bright spots of the reference. These spots may come from residual spatial drift across the *b* = 0 repeats from which the reference was estimated. Figure 3b–c quantify agreement with the signal and the noise reference, with the proposed method matching both better than MPPCA. In DTI and DKI metrics, denoising improves agreement with the reference, with neither method clearly ahead (see Figure 3d–e). Figure S4 shows the DTI and DKI maps for the slice of Figure 3a.

**Figure 3:**
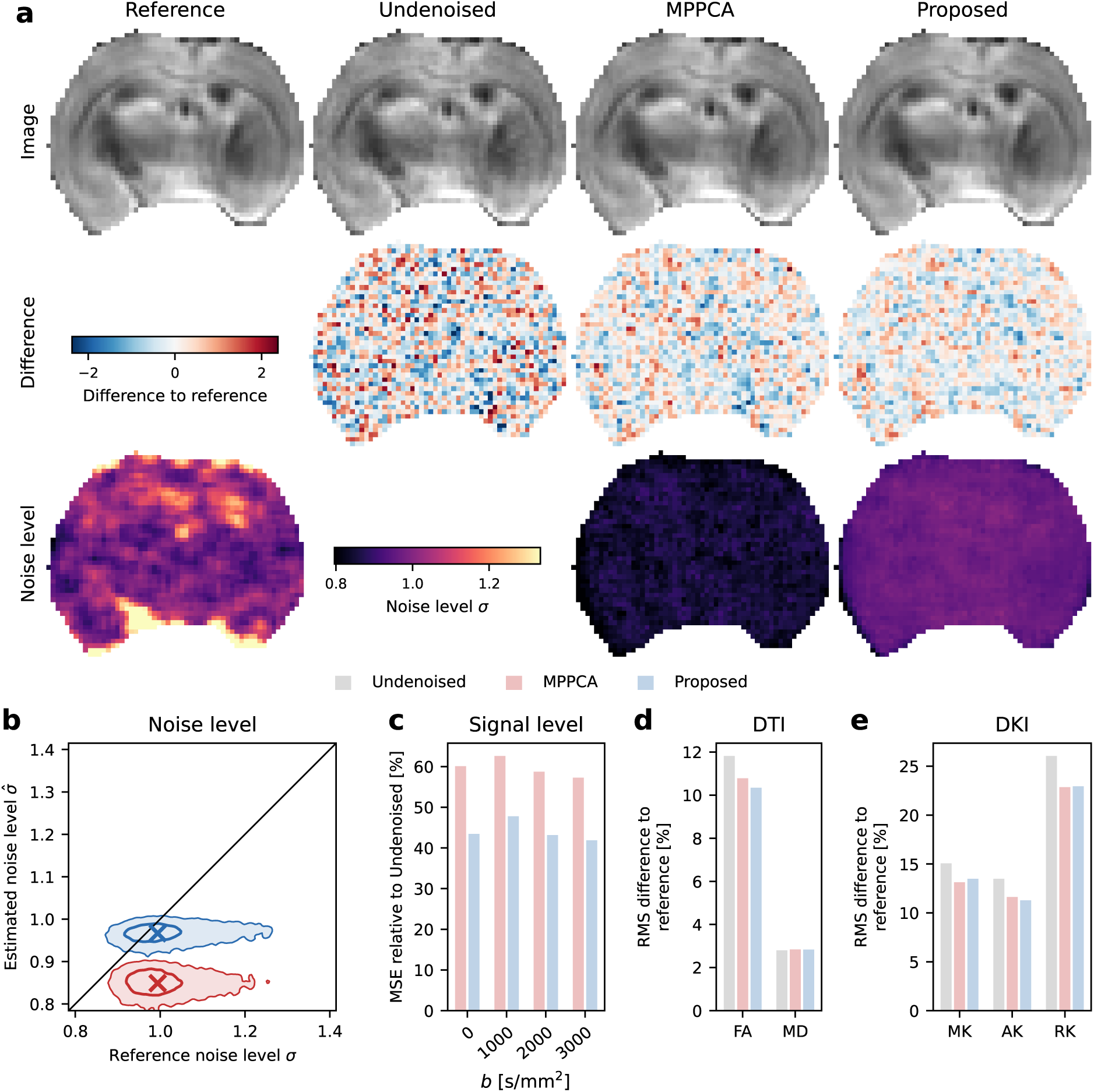
Ex vivo mouse brain dMRI. **(a)** Qualitative comparison of denoising methods against the reference. Rows show a diffusion-weighted slice at *b* = 2000 s*/*mm^2^, the difference relative to the reference, and the noise level *σ*. **(b)** Voxel-wise comparison of the estimated noise level *σ*^ against the reference *σ*. Crosses mark the medians, contours the central 50 % and 90 % of the voxels, and the diagonal unity. **(c)** Signal mean squared error (MSE) per shell, as a percentage of the undenoised series’ MSE. **(d–e)** Root mean square error in DTI and DKI metric against the reference, normalized to the reference metric’s mean over white matter. DTI metrics include fractional anisotropy (FA) and mean diffusivity (MD); DKI metrics include mean, axial and radial kurtosis (MK, AK and RK). Figure S4 shows maps for the slice in (a).

### 3.3 In vivo consistency within and across scanners

We evaluated the proposed method on human in vivo data (Manzano Patron et al., 2024, 2026; Warring-ton et al., 2023, 2025b), offering repeated scans on one scanner to assess repeatability and scans across multiple scanners to assess reproducibility.

Figure 4 addresses repeatability at 0.9 mm isotropic resolution on magnitude dMRI, with one subject scanned in four repeats. Denoising visibly reduces the noise, but the outputs of the two denoising methods differ (see Figure 4a). The proposed method removes a pronounced grainy texture that MPPCA keeps, which amounts to more aggressive denoising. The more aggressive denoising follows from a higher estimated noise level (see Figure 4b). The proposed method also yields the lowest signal mean squared error (MSE) against the other repeats’ mean (see Figure 4c). The MSE reduction is driven by lower variance, at a slight increase in bias. In both DTI and DKI metrics, the proposed method lowers the coefficient of variation (CoV) across repeats (see Figure 4d–e). Figure S5 shows the FA and MK maps behind this analysis, and Figures S6 and S7 repeat the analysis and the maps on complex data, with NORDIC as a further comparison.

**Figure 4:**
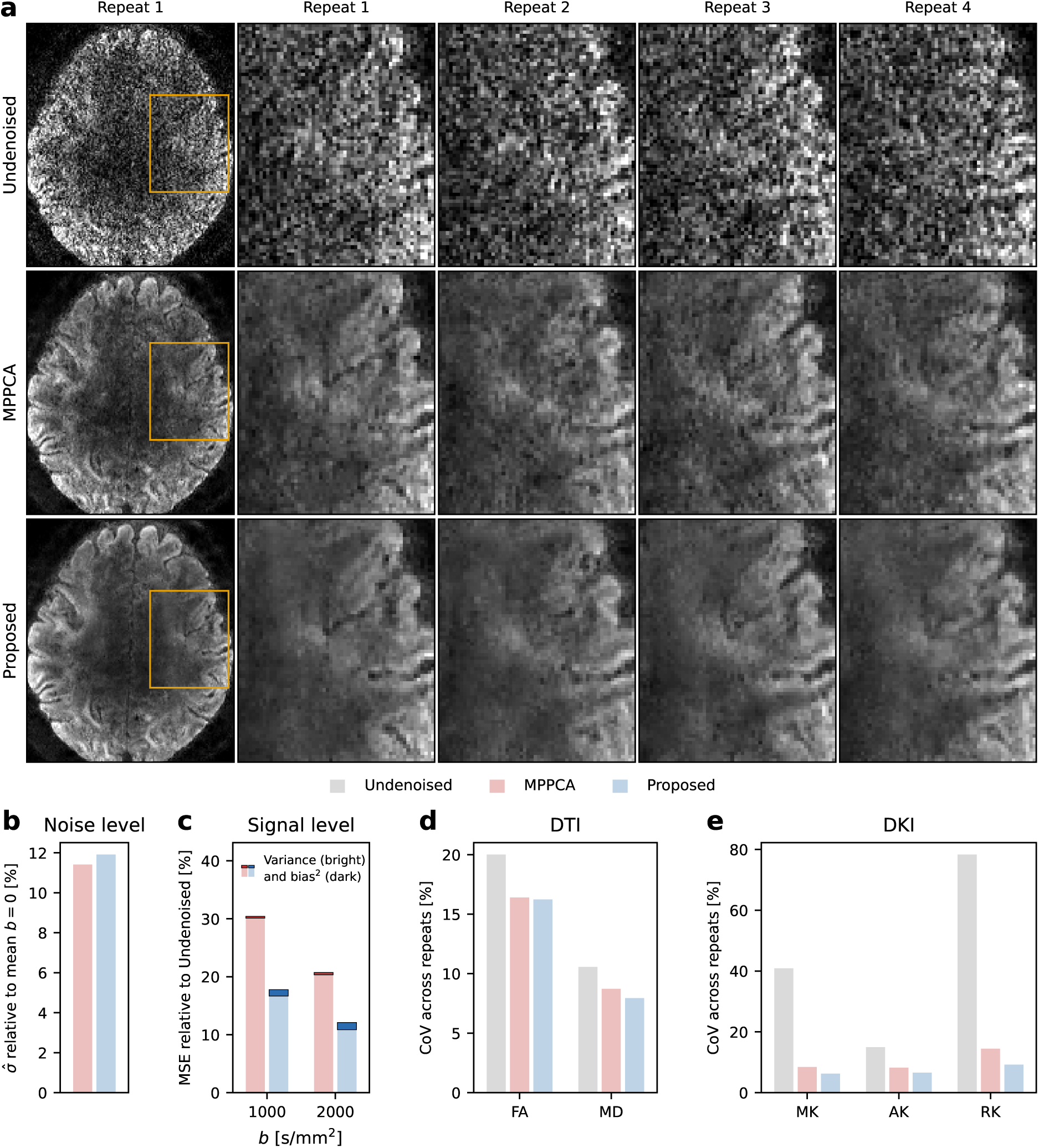
In vivo repeatability within one scanner at 0.9 mm. (**a**) Qualitative comparison of the undenoised series and the denoising methods for one axial slice at *b* = 2000 s*/*mm^2^. Columns show four repeats of one subject on one scanner at 0.9 mm isotropic resolution, magnified on the region highlighted in the leading column. **(b–e)** Quantitative comparison across the four repeats. **(b)** Noise level *σ*^ estimated by each denoising method. **(c)** Per-shell signal mean squared error (MSE) against the other repeats’ mean, split into variance and squared bias. **(d–e)** Coefficient of variation (CoV) for DTI and DKI metrics across the four repeats. DTI includes fractional anisotropy (FA) and mean diffusivity (MD); DKI includes mean, axial and radial kurtosis (MK, AK and RK). Through (b–e), data are pooled over all voxels of the brain (b–c) or white matter (d–e). Bars give the median across the four repeats in (b) and the mean in (c). Figure S5 shows FA and MK maps for the slice in (a).

Figure 5 further examines repeatability across a broader range of seven scanners at 2 mm isotropic resolution on magnitude dMRI, with one subject per scanner scanned in six repeats. Compared to Figure 4a, the differences between undenoised and denoised images, as well as among denoising methods, are much less pronounced (see Figure 5a). Nonetheless, the proposed method removes extra spurious texture that MPPCA keeps (red arrows). The proposed method again reads a higher noise level than MPPCA, and the correction relative to that level is no smaller than in Figure 4b (see Figure 5b). However, the absolute noise level in Figure 5b is much lower, so the correction moves far less noise in total, explaining the modest visible differences in Figure 5a. Similarly, the proposed method reduces signal MSE and DKI metric CoV over MPPCA, less than in Figure 4 but on every scanner, with DTI metrics on par (see Figure 5c–e). Figure S8 shows the FA and MK maps behind this analysis.

**Figure 5:**
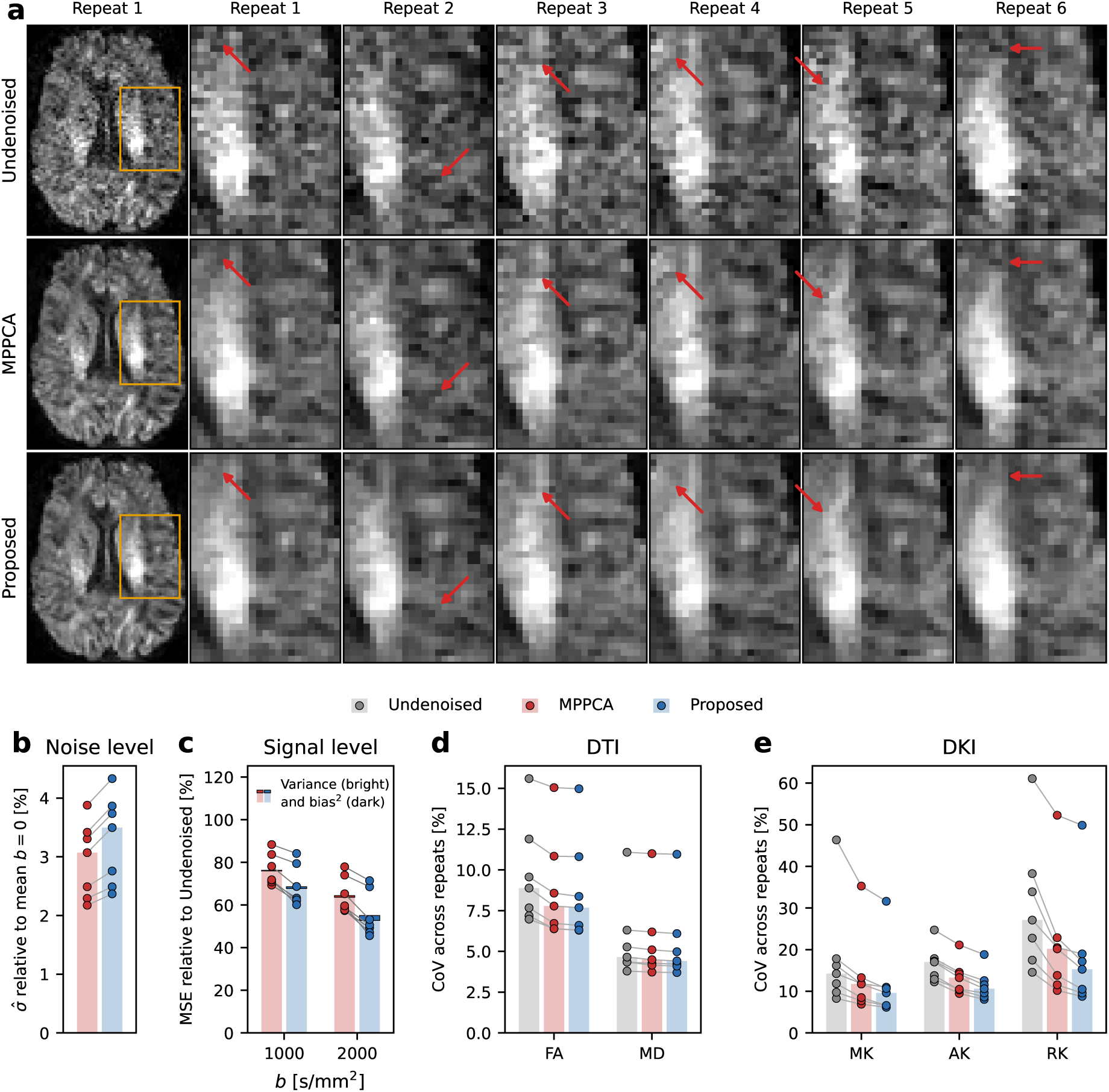
In vivo repeatability within scanners at 2 mm. (**a**) Qualitative comparison of the undenoised series and the denoising methods for one axial slice at *b* = 2000 s*/*mm^2^. Columns show the six repeats of one subject on a Philips Ingenia with a 32-channel coil, magnified on the region highlighted in the leading column. Red arrows mark matching locations across rows. **(b–e)** Quantitative comparison across seven scanners with six repeats each. **(b)** Noise level *σ*^ estimated by each denoising method. **(c)** Per-shell signal mean squared error (MSE) against the other repeats’ mean, split into variance and squared bias. **(d–e)** Coefficient of variation (CoV) for DTI and DKI metrics across repeats. DTI includes fractional anisotropy (FA) and mean diffusivity (MD); DKI includes mean, axial and radial kurtosis (MK, AK and RK). Through (b–e), each point denotes one scanner, with data pooled over the six repeats and all voxels of the brain (b–c) or white matter (d–e). Bars give the median across scanners, except in (c), where the bar is the mean. Figure S8 shows FA and MK maps for the slice in (a).

Figure 6 examines reproducibility across the same seven scanners as in Figure 5, but with seven subjects each scanned once on five or six of the scanners. Images differ notably between scanners, more than the undenoised input differs from either denoised series (see Figure 6a). While denoised outputs look very similar, the proposed method removes spurious textures that MPPCA keeps, more pronounced on Philips and GE scanners (red arrows). The estimated noise levels match those of Figure 5b, reflecting that acquisitions are shared and only subjects differ (see Figure 6b). In DTI metrics, the two denoising methods are on par, whereas in DKI metrics the proposed method lowers the CoV across scanners (see Figure 6c–d). Compared to within-scanner values, the CoV is moderately inflated, which indicates differences between the scanners and their image reconstructions. Figure S10 shows the FA and MK maps behind this analysis.

**Figure 6:**
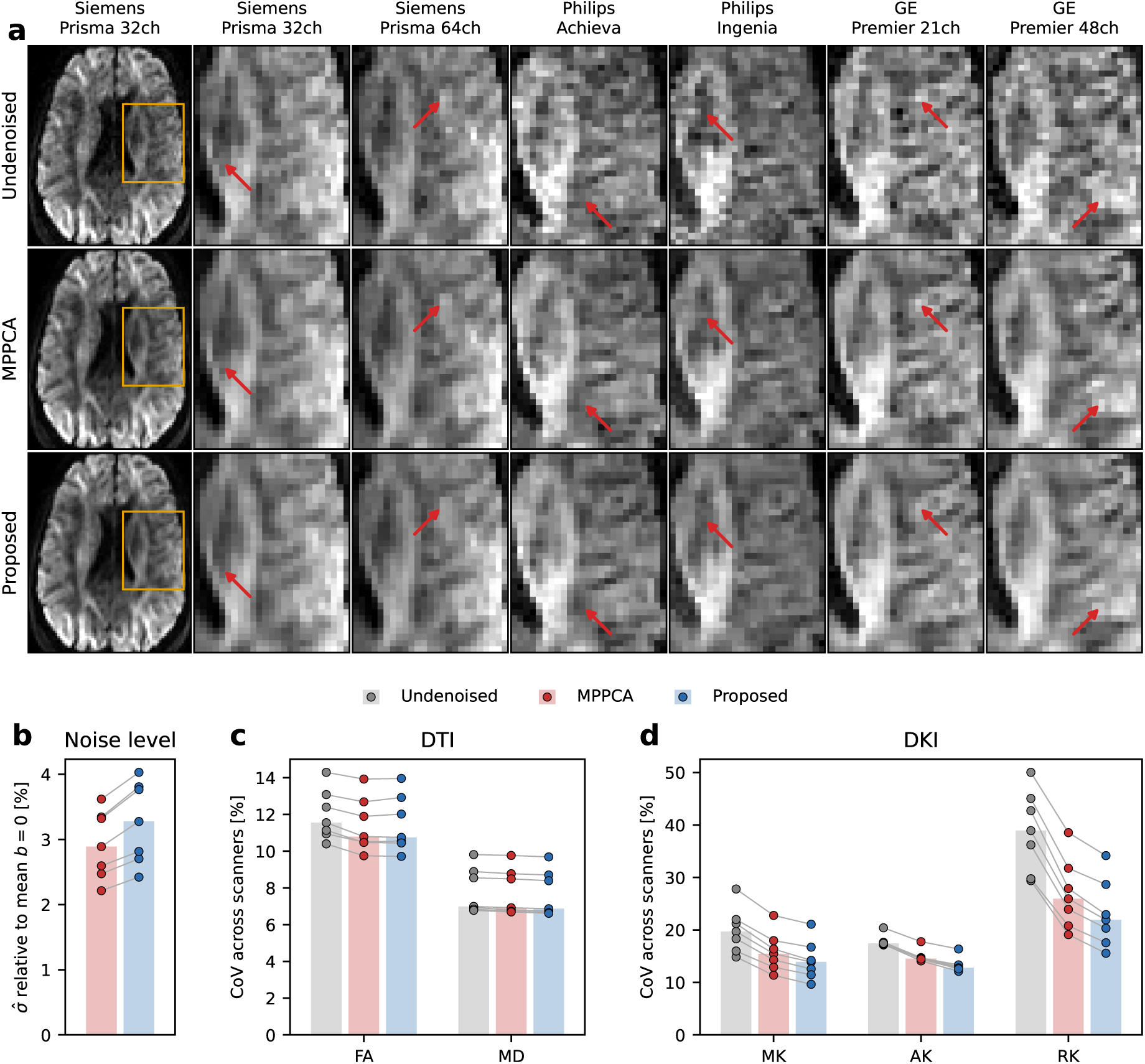
In vivo reproducibility across scanners at 2 mm. (**a**) Qualitative comparison of denoising methods for one axial slice at *b* = 2000 s*/*mm^2^. Columns show the six scanners of one subject’s cohort, magnified on the region highlighted in the leading column. Red arrows mark matching locations across rows. **(b–d)** Quantitative comparison across seven subjects, each scanned once on five or six of seven scanners. **(b)** Noise level *σ*^ estimated by each denoising method. Each point denotes one scanner, averaged over the subjects scanned there, and bars denote the median across scanners. **(c–d)** Coefficient of variation (CoV) for DTI and DKI metrics across scanners. DTI includes fractional anisotropy (FA) and mean diffusivity (MD); DKI includes mean, axial and radial kurtosis (MK, AK and RK). Each point denotes one subject and bars the median across subjects, with data pooled over all white matter voxels. Figure S10 shows FA and MK maps for the slice in (a).

### 3.4 The noise covariance across public datasets

We applied the proposed method to 28 public datasets, which let us characterize the noise covariance across a wide range of scanners, acquisitions and reconstructions.

Figure 7a shows the noise level correction the proposed method applies to MPPCA’s estimate on its horizontal axis, with the distribution across the datasets above it. The correction is substantial, with a median of 8.2 %, and positive throughout. Thus, if the estimated noise covariance holds, MPPCA reads the noise level too low and removes too little noise in every scan. The correction size likely reflects the scanner and its image reconstruction. Scan-dependent susceptibility, eddy currents, motion or other effects may add to it, which Figure S9 investigates on the repeat scans of Figure 5. There is no reference for the noise level in vivo, but the estimated noise covariance predicts its impact on MPPCA’s reading of the noise level accurately, with a median prediction error of 1.2 % on the vertical axis. Figure 7b attributes the applied noise level correction to the two components of the noise covariance model, the spatial variation of the noise level and the spatial noise correlation. The correlation alone applies about 99 % of the full model’s correction, while the spatial variation alone applies only about 2 %.

**Figure 7:**
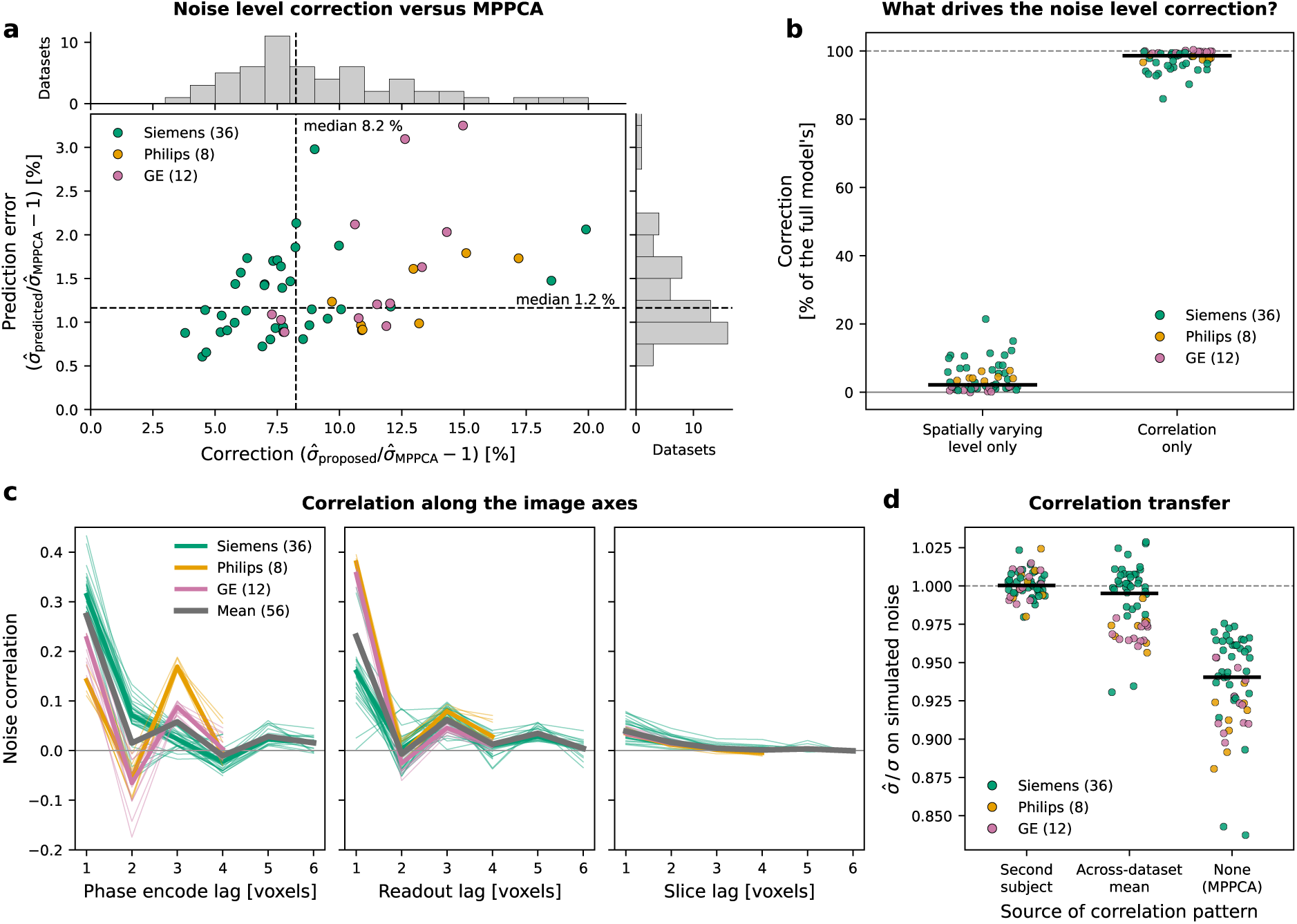
The noise covariance across public datasets. **(a)** Error of the noise level predicted for MPPCA from the estimated noise covariance, against the noise level correction the proposed method applies relative to MPPCA. Each point denotes one of 56 scans from 28 public datasets with two subjects each, colored by scanner vendor. Histograms give the distribution along each axis and dashed lines their medians. **(b)** Attribution of the correction in (a) to the two components of the noise covariance model, the spatially varying level and the correlation, each as a share of the full model’s. **(c)** Noise correlation against the lag along the phase encode, readout and slice axes, one line per scan with the vendor medians and the mean over all scans drawn thick. The curves give one side of the symmetric correlation, corresponding to the central row and column of the 2D correlation pattern in Figure 1. The lag support differs between scans due to different patch sizes. **(d)** Recovery of a known noise level on synthesized noise, after whitening with a correlation from elsewhere. The correlation is taken from the second subject of the same dataset, from the mean over the other datasets, or not applied at all (MPPCA).

Figure 7c shows that the correlation carries vendor-specific patterns along phase encode and readout directions, likely due to partial Fourier, regridding and in-plane filtering. Partial Fourier and the in-plane acceleration technique largely coincide with the vendor across our datasets (see Table A2), so the two cannot be separated here. In contrast, the slice direction carries little correlation, as expected for in-plane reconstruction. Further, Figure 7d studies whether the correlation transfers across subjects. Indeed, the proposed method recovers the level the noise was synthesized with, even when whitening with a second subject’s correlation from the same dataset. In contrast, an average correlation pattern, computed over the other datasets, scatters widely around the true noise level. When not accounting for correlation, as in MPPCA, the noise level is underestimated, but less than on the real data in Figure 7a, where signal is present in addition to the noise. Importantly, Figure S11 shows that the correlation also transfers across brain regions, further supporting stationarity of the estimated correlation patterns.

## 4 Discussion

Marčenko–Pastur (MP) denoising is widely used in diffusion magnetic resonance imaging (dMRI) processing, but the i.i.d. noise it assumes is not what most reconstruction pipelines deliver. We have proposed an extension that restores the i.i.d. noise assumption, without additional measurements or knowledge of the reconstruction. Compared to the widely used MPPCA method (Veraart et al., 2016c), the proposed method improves agreement with repeated ex vivo and in vivo measurements in signal and noise level, and yields more consistent diffusion metrics. Application to a broad panel of 28 public datasets revealed a noise covariance footprint in every one of them, affecting MP denoising mainly through correlation between neighboring voxels rather than through a spatially varying noise level.

### Common reconstructions introduce correlated noise worth accounting for

We found reconstruction-induced deviations from the i.i.d. noise assumption in every dataset we examined, across vendors, acquisitions and reconstructions (see Figure 7a). For MPPCA (Veraart et al., 2016c), these deviations leave the estimated noise level about 8 % too low, so part of the noise is kept as signal. This number may well understate the problem, as our results suggest the deviations are largely correlated noise (see Figure 7b). What is kept under correlated noise is not the per-voxel grain that an i.i.d. assumption would predict, but fine structure that is easily confused with anatomical detail, by the eye, by the denoiser, and potentially by downstream microstructure analyses. In contrast, the spatial variation of the noise level had a negligible effect throughout the data investigated, suggesting that typical within-patch variation is smooth enough to approximately satisfy the assumption of identically distributed noise. Yet, our estimate of the spatially varying noise level is smooth by construction and follows gradual variation only, which may fall short in data with very strong spatial noise level variation (Cordero-Grande et al., 2019; Moeller et al., 2021).

### The noise correlation tracks the vendor reconstruction

Generalizing across reconstructions cannot come without simplifying assumptions. Our core assumption over previous work (Henriques et al., 2023; Moeller et al., 2021) is that the within-patch noise correlation is stationary across the brain. Indeed, we show that estimated correlations transfer across brain regions (see Figure S11). Yet, a reconstruction-induced correlation would imply more than this assumption, and our results are in line with it. The estimated correlation patterns act in-plane, are vendor-specific, and transfer across subjects (see Figure 7c–d). We expect few effects behind the short-range correlation relevant to patch-based denoising, essentially partial Fourier and the readout. While this invites explicit modeling, correlation patterns arise from often-undisclosed vendor implementations (D’Antonio et al., 2026), so hand-built models may only hold until the next software release.

### MP denoising restored, not overdone

Across simulations, ex vivo and in vivo data, the proposed method lowered the signal mean squared error against the reference. The improvement came from reduced variance at a slight increase in bias (see Figure 2f–g, Figure 4c and Figure 5c). Our results suggest this bias is the one MP denoising carries by design, rather than a sign of denoising more aggressively. In simulations, both methods carried a similar bias in the absence of correlation, the setting MPPCA assumes. Under correlation, MPPCA shed that bias and took on substantially more noise variance instead, an unintended trade that follows from fitting its threshold under a broken assumption. The proposed method’s improved placement of the split between signal and noise is further supported by the ex vivo reference, where it estimated the noise level more accurately (see Figure 3b).

### Reduced noise variance improves diffusion kurtosis imaging consistency

The impact of denoising on diffusion metrics followed the signal-to-noise ratio (SNR) of the data and the SNR demand of the metric (Henriques et al., 2021; Jensen et al., 2005). At the relatively high SNR of the ex vivo data, neither diffusion tensor imaging (DTI) nor diffusion kurtosis imaging (DKI) accuracy against the reference differed much across denoising methods (see Figure 3d–e). Across in vivo repeats, the proposed method yielded more consistent DKI metrics than MPPCA (see Figure 4e and Figure 5e). In contrast, DTI metrics improved only at the lower SNR of the 0.9 mm dataset (see Figure 4d). The observations on DTI and DKI consistency carried over to comparisons across scanners (see Figure 6c–d). There, denoising can only reduce the measurement noise, while systematic scanner differences remain (Warrington et al., 2023).

### One noise covariance, but its impact may vary

What a given noise covariance costs in MP denoising is also shaped by the signal and by the acquisition. MP denoising assumes that the diffusion signal is carried by few components, well separated from the noise. However, distortion and motion across volumes add weak components, blurring the separation between signal and noise. Indeed, we found an association between such effects and the impact of the noise covariance on denoising across repeated scans (see Figure S9), in line with previous test-retest analyses on the same data (Warrington et al., 2023). Therefore, restoring MP’s assumption on the signal may deserve more attention, while advances have focused almost exclusively on the noise (Cordero-Grande et al., 2019; Manzano Patron et al., 2024; Moeller et al., 2021; Ye et al., 2025).

Further, how much the noise covariance affects MP denoising depends on the number of volumes relative to the patch size (see Figure S3). This may lead to unexpected results when comparing otherwise identical acquisitions that differ in the number of volumes. The proposed method largely removes this dependency.

### The proposed method in the landscape of MP denoising

Our main contribution is to make MP denoising compatible with non-i.i.d. noise. Existing methods account only for a spatially varying noise level (Moeller et al., 2021), require additional measurements (Cordero-Grande et al., 2019; Henriques et al., 2023; Moeller et al., 2021) or knowledge of the reconstruction (Cordero-Grande et al., 2019; Moeller et al., 2021). The proposed method needs neither additional measurements nor knowledge of the reconstruction, and accounts for a spatially varying noise level and for correlation within a patch. We estimate both the noise level and the threshold from the diffusion series itself, similar to MPPCA (Veraart et al., 2016c) and to a NORDIC variant designed for complex dMRI (Moeller, 2026; Moeller et al., 2021). We therefore benchmarked the proposed method against MPPCA, the widely used choice on magnitude dMRI. Under the same patch size rule, the noise covariance model is then the only difference between the two. The proposed method also applies to complex dMRI, where we additionally compared against NORDIC (see Figures S6 and S7). The method like-wise combines with downstream variants of the denoising step, such as patch averaging (Manjón et al., 2013), eigenvalue shrinkage (Gavish & Donoho, 2017; Leeb & Romanov, 2021), alternative threshold rules (Henriques et al., 2023; Olesen et al., 2023) and adaptive patch selection (Ades-Aron et al., 2025).

### Limitations and future directions

Our reported noise level excludes the flat mode, the one direction the estimator can only weakly deter-mine, and may therefore fall below the voxel’s full noise level. However, our simulations and ex vivo data suggest this potential bias is much smaller than the one MPPCA carries under the broken i.i.d. noise assumption (see Figure 2d–e and Figure 3b). The improved noise level estimate may be useful for several downstream applications, such as parameter estimation under noise (Ades-Aron et al., 2018; David et al., 2024; Koay & Basser, 2006; Maximov et al., 2019; Mezzano et al., 2026; París et al., 2026), uncertainty quantification (Harms et al., 2024; Harms & Roebroeck, 2018), and training data synthesis (Ioakeimidis et al., 2026). For the latter, the estimated correlation may make synthesized noise more realistic than the level alone (Xue et al., 2025).

The proposed noise covariance model assumes the correlation to be stationary across the brain, and the noise to be additive with a fixed covariance. Both assumptions can be broken, for example by non-Cartesian trajectories (Lee et al., 2023), learned reconstructions (D’Antonio et al., 2026) or resampling during processing (Rohde et al., 2005).

Due to its public availability across a wide range of scanners, acquisitions and reconstructions, we focused on magnitude dMRI. Denoising magnitude dMRI is considered suboptimal, because the Rician noise floor breaks the additive noise model (Eichner et al., 2015; Gudbjartsson & Patz, 1995), yet it remains effective in practice (Schilling et al., 2026). We also applied the proposed method to complex dMRI, using one of various approaches. We removed the background phase (Moeller, 2026; Sprenger et al., 2017) and denoised the real part after rotation onto the real axis (Eichner et al., 2015) (see Figures S6 and S7). To improve complex denoising with the proposed method, future work would have to extend the correlation model and assess the phase handling, which itself shapes the noise covariance (Sprenger et al., 2017).

Beyond dMRI, the proposed method should apply to other multi-volume series with a sufficient number of volumes (see Figure S1a), such as functional MRI (Vizioli et al., 2021) or multi-parameter mapping (Tabelow et al., 2019).

## 5 Conclusion

Marčenko–Pastur denoising is widely used in diffusion magnetic resonance imaging, but under-denoises once image reconstruction has reshaped the noise. We restore the intended denoising post hoc, without additional measurements or knowledge of the reconstruction. The method therefore applies to existing and new data alike, opening headroom for microstructure analyses.

## Data and Code Availability

All data analyzed in this study are publicly available. The subject phantom and the 0.9 mm repeat data are from Manzano Patron et al., 2024, 2026, the ex vivo mouse brain data from Henriques et al., 2023, and the ON-Harmony data from Warrington et al., 2023, 2025b. The remaining datasets were obtained from OpenNeuro.org and the Human Connectome Project and are listed in Table A2.

An implementation of the denoising algorithm is available at https://github.com/quantitative-mri-and-in-vivo-histology/camppca (commit b93388b), and the code to reproduce the results of this paper at https://github.com/quantitative-mri-and-in-vivo-histology/camppca_paper (commit d050600).

## Author Contributions

L.M.: conceptualization, methodology, software, data curation, investigation, formal analysis, valida-tion, visualization, writing of the original draft. K.T.: review and editing. S.M.: funding acquisition, supervision, review and editing.

## Funding

This work was supported by the German Research Foundation (DFG Priority Program 2041 “Computational Connectomics”, [MO 2397/5-1, MO 2397/5-2], by the Emmy Noether Stipend: MO 2397/4-1; MO 2397/4-2).

Funded by the European Union. Views and opinions expressed are however those of the author(s) only and do not necessarily reflect those of the European Union or the European Research Council Executive Agency. Neither the European Union nor the granting authority can be held responsible for them. This work is supported by ERC grant (Acronym: MRStain, Grant agreement ID: 101089218, DOI: 10.3030/101089218).

## Ethics

This study did not acquire new data. Ethical approval and informed consent of the human participants, and approval of the animal procedures for the ex vivo mouse brain data, were obtained by the original studies of the datasets listed in Data and Code Availability.

## Declaration of Competing Interests

The authors declare no competing interests.

## Declaration of the use of AI

The authors used Anthropic Claude Opus to assist with drafting and editing manuscript text and with developing the software used for the analyses and figures. All figures were produced by that code from the measured data; no figure content was generated by AI. All content was reviewed and verified by the authors, who take full responsibility for the manuscript.

## A Dataset information

**Table A1:** Dataset information: In vivo consistency within and across scanners. One row per scanner, in two blocks. The ON-Harmony block (Warrington et al., 2023, 2025b) holds a repeat subject scanned six times on that scanner, while every other subject of the same study phase contributed one scan, and each shell holds 50 directions. The EDDEN block (Manzano Patron et al., 2024, 2026) holds one subject scanned in four repeats, available as complex data. Remaining volumes are *b* = 0. All series were acquired with anterior-posterior phase encoding, with a reversed-polarity *b* = 0 series beside them. Symbols: – the factor was not used.

| Scanner | Repeat subject | Coil (ch) | Voxel (mm) | *b* (s/mm<sup>2</sup>) | Vols. | PF | *R* | MB | TE/TR (ms) |
|---|---|---|---|---|---|---|---|---|---|
| *ON-Harmony* |  |  |  |  |  |  |  |  |  |
| Siemens Prisma | sub-13192 | 32 | 2.02 | 0/1000/2000 | 105 | 6/8 | – | 3 | 92/3600 |
| Siemens Prisma | sub-14229 | 64 | 2.02 | 0/1000/2000 | 105 | 6/8 | – | 3 | 92/3600 |
| Siemens Trio | sub-14230 | 32 | 2.02 | 0/1000/2000 | 105 | 6/8 | – | 3 | 96/3600 |
| Philips Achieva | sub-14482 | 32 | 2.00 | 0/1000/2000 | 107 | 4/5 | 2 | – | 71/10400 |
| Philips Ingenia | sub-15320 | 32 | 2.00 | 0/1000/2000 | 105 | 4/5 | 1.5 | 3 | 98/4370 |
| GE Premier | sub-16745 | 48 | 2.02 | 0/1000/2000 | 106 | – | 2 | 2 | 68/4900 |
| GE Premier | sub-16975 | 21 | 2.02 | 0/1000/2000 | 106 | – | 2 | 2 | 69/8000 |
| *EDDEN* |  |  |  |  |  |  |  |  |  |
| Siemens Prisma | sub-01 | 32 | 0.90 | 0/1000/2000 | 199 | 6/8 | 2 | 3 | 91/6569 |

**Table A2:** Dataset information: The noise covariance across public datasets. Each row contains a dataset with two subjects, grouped by release. Each subject contributes one scan, and no subject appears in more than one dataset. Symbols: – the factor was not used or does not apply,the value could not be inferred, ^a^ the two subjects were acquired at different echo times, ^b^ *b*-values sampled over a continuous range rather than on shells.

| Reference | Subjects | Scanner | *B*<sub>0</sub><br>(T) | Coil<br>(ch) | Voxel<br>(mm) | *b*<br>(s/mm<sup>2</sup>) | Vols. | PF | In-plane<br>technique | *R* | MB | TE/TR<br>(ms) |
|---|---|---|---|---|---|---|---|---|---|---|---|---|
| Zhang et al., 2025 | sub-001, sub-002 | Siemens Prisma fit | 3 | 64 | 2.00 | 0/1000/2000 | 141 | 6/8 | - | - | - | 82/9190 |
|  | sub-003, sub-004 | Siemens Prisma fit | 3 | 64 | 2.00 | 0/1000/2000 | 141 | 6/8 | - | - | 3 | 93/3378 |
|  | sub-005, sub-006 | Siemens Prisma fit | 3 | 64 | 2.00 | 0/1000/2000 | 141 | 6/8 | k-space | 2 | 3 | 82/2740 |
|  | sub-801, sub-803 | Siemens Prisma fit | 3 | 64 | 2.00 | 0/1000/2000 | 141 | 6/8 | - | - | 6 | 93/1723 |
|  | sub-806, sub-808 | Siemens Prisma fit | 3 | 64 | 2.00 | 0/1000/2000 | 141 | 6/8 | k-space | 2 | 6 | 82/1404 |
| Figueroa-Vargas et al., 2024 | sub-001, sub-003 | Siemens Skyra | 3 | ? | 1.88 | 0/1000 | 93 | ? | k-space | 2 | - | 95/8600 |
| Bhatt et al., 2026 | sub-02, sub-03 | Siemens Magnetom Connectom.X | 3 | 72 | 2.00 | 0/50/350/800/1500 | 150 | 6/8 | k-space | 2 | 2 | 53/3800 |
| Way et al., 2025 | sub-s001, sub-s002 | Siemens Prisma | 3 | ? | 2.00 | 0/2000 | 65 | ? | - | - | 2 | 85/4570 |
| Van Essen et al., 2013 | 100206, 103818 | Siemens Connectome Skyra | 3 | 32 | 1.25 | 0/1000/2000/3000 | 288 | 6/8 | - | - | 3 | 90/5520 |
| Van Essen et al., 2013 | 104416, 114823 | Siemens Magnetom 7T | 7 | 32 | 1.05 | 0/1000/2000 | 143 | 6/8 | k-space | 3 | 2 | 71/7000 |
| Taylor et al., 2026 | sub-1, sub-103 | GE Discovery MR750 | 3 | 32 | 2.00 | 0/300/700/2500 | 115 | ? | - | - | - | 74/7600 |
| Babayan et al., 2020 | sub-010002, sub-010003 | Siemens Verio | 3 | 32 | 1.72 | 0/1000 | 67 | 7/8 | k-space | 2 | 2 | 80/7000 |
| Velasco, 2026 | sub-wlsubj016, sub-wlsubj119 | Siemens Prisma | 3 | 64 | 1.50 | 0/1500 | 138 | 6/8 | - | - | 3 | 85/4150 |
| Warrington et al., 2025a | sub-16842, sub-16974 | Siemens Prisma | 3 | 32 | 2.02 | 0/1000/2000 | 105 | 6/8 | - | - | 3 | 92/3600 |
|  | sub-16975, sub-16981 | Siemens Prisma | 3 | 64 | 2.02 | 0/1000/2000 | 105 | 6/8 | - | - | 3 | 92/3600 |
|  | sub-10975, sub-12813 | Siemens TrioTim | 3 | 32 | 2.02 | 0/1000/2000 | 105 | 6/8 | - | - | 3 | 96/3600 |
|  | sub-14482, sub-16841 | Philips Achieva | 3 | 32 | 2.00 | 0/1000/2000 | 107 | 4/5 | image dom. | 2 | - | 71/10400 |
|  | sub-13192, sub-13305 | Philips Achieva | 3 | 32 | 2.00 | 0/1000/2000 | 105 | 4/5 | image dom. | 2 | - | 70/9900 |
|  | sub-14221, sub-16794 | Philips Ingenia | 3 | 32 | 2.00 | 0/1000/2000 | 105 | 4/5 | image dom. | 1.5 | 3 | 98/4370 |
|  | sub-14229, sub-14230 | Philips Ingenia | 3 | 32 | 2.00 | 0/1000/2000 | 105 | 4/5 | image dom. | 2 | 3 | 89/4020 |
|  | sub-03286, sub-03997 | GE Discovery MR750 | 3 | 32 | 2.02 | 0/1000 | 65 | ? | ? | 2 | - | 65,72<sup>a</sup>/11000 |
|  | sub-16745, sub-16793 | GE SIGNA Premier | 3 | 48 | 2.02 | 0/1000/2000 | 106 | ? | ? | 2 | 2 | 68/4900 |
|  | sub-15320, sub-16766 | GE SIGNA Premier | 3 | 21 | 2.02 | 0/1000/2000 | 106 | ? | ? | 2 | 2 | 70/8000 |
| Hau et al., 2025 | sub-ts003, sub-ts005 | GE Discovery MR750 | 3 | 8 | 1.88 | 0/1000 | 62 | ? | image dom. | 2 | - | 82/8500 |
|  | sub-ts007, sub-ts008 | Siemens Prisma | 3 | 32 | 1.50 | 0/1500/3000 | 199 | 6/8 | - | - | 4 | 89/3230 |
| Pavon et al., 2022 | sub-CON015, sub-CON025 | Siemens Prisma | 3 | 64 | 2.00 | 0/150–4000<sup>b</sup> | 272 | 6/8 | k-space | 2 | 3 | 102/4000 |
| Coutiño et al., 2026 | sub-15464, sub-700 | GE Discovery MR750 | 3 | 32 | 1.72 | 0/800/1500/2500 | 121 | ? | image dom. | 2 | 2 | 81/4000 |
| Shaw et al., 2026 | sub-H0101, sub-H0102 | Siemens Magnetom 7T | 7 | 32 | 1.80 | 0/1000/2500 | 96 | 6/8 | k-space | 3 | - | 64/8000 |

## Supplementary Material

**Figure S1:**
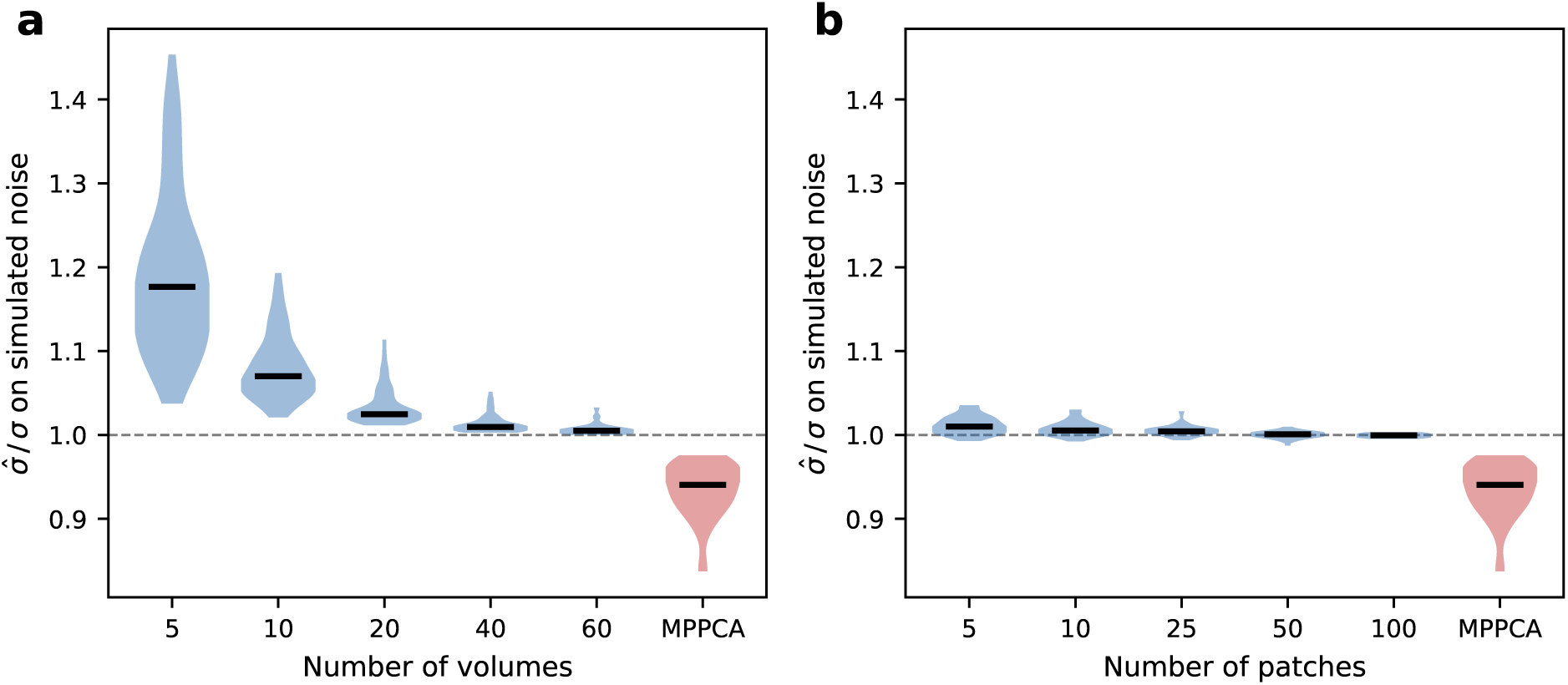
Volumes and patches needed to estimate the correlation. Recovery of a known noise level on synthesized noise, after whitening with a correlation estimated from **(a)** a subset of volumes and **(b)** a subset of patches. Violins span the values with the median marked, and the dashed line sits at the known noise level. The last violin of each panel leaves the noise unwhitened for reference, as in MPPCA. The noise is synthesized with the correlation estimated on all volumes and all patches, for each of 56 scans from 28 public datasets with two subjects each (see Section 2.6). Subsets of volumes keep each *b*-shell’s share, three per scan, against one subset of patches per scan. The number of volumes influences the estimated correlation more than the number of patches.

**Figure S2:**
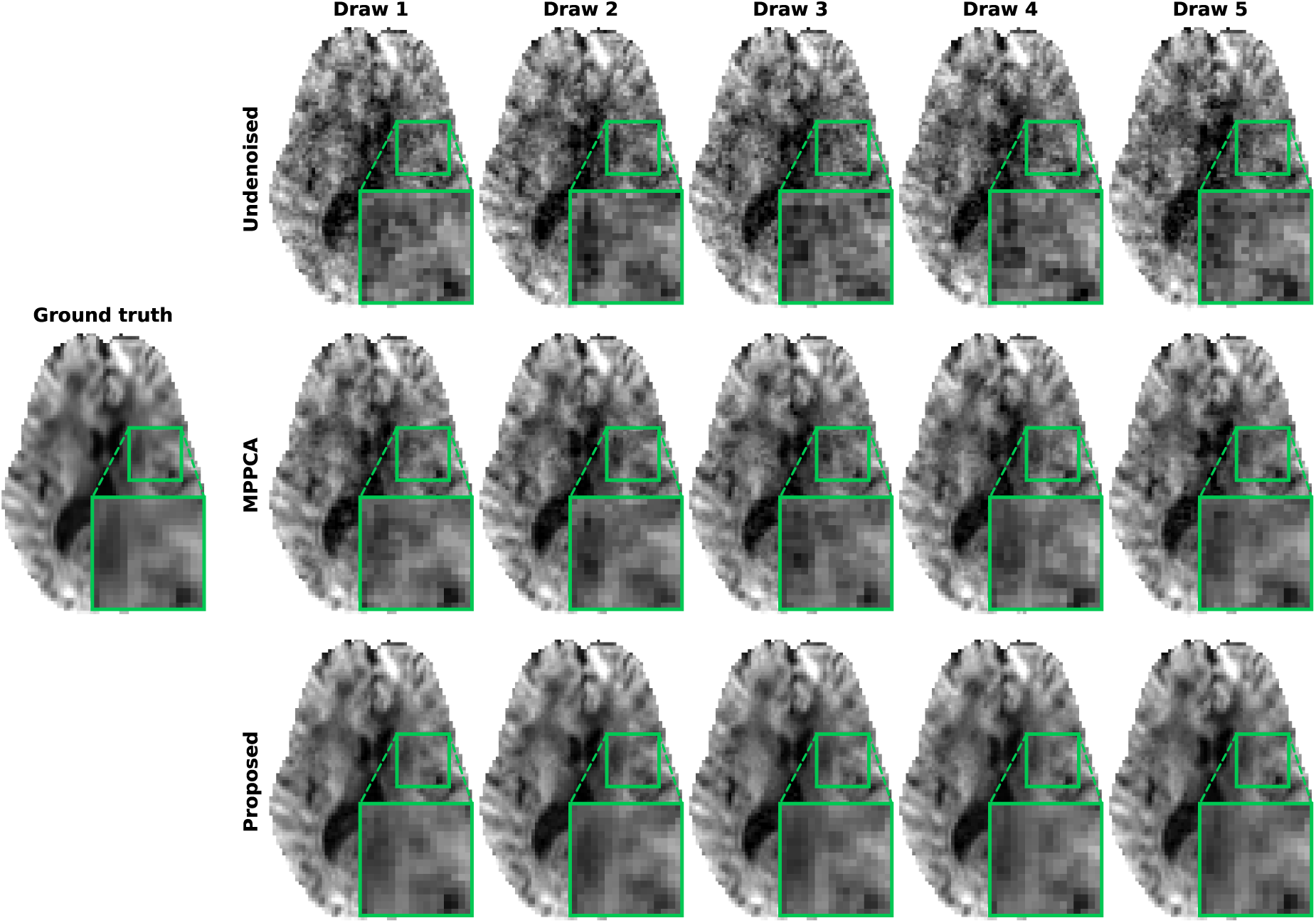
Simulation: qualitative comparison across independent noise draws. One diffusion-weighted slice at *b* = 2000 s*/*mm^2^, with the noise-free ground truth on the left and a comparison across denoising methods in the rows to its right. Columns show five independent noise draws of the setting in Figure 2a, illustrating how noise can shape anatomy-like structure, varying across noise draws. Insets magnify the outlined region. While structures vary across draws under both denoising methods, the proposed method gives a more consistent appearance.

**Figure S3:**
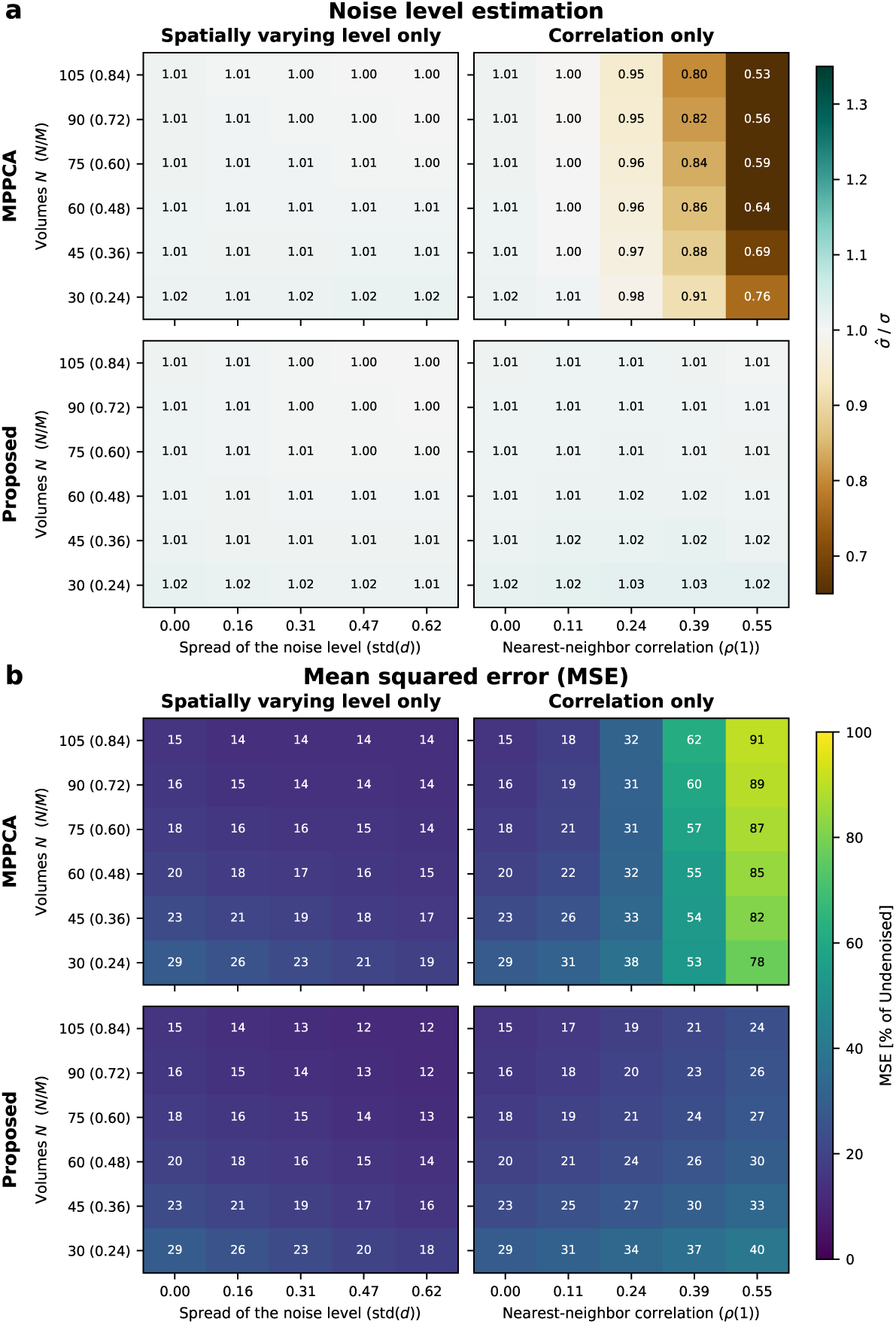
Simulation: the noise covariance across matrix aspect ratios. **(a)** Estimated noise level (*σ*^) relative to the ground truth noise level (*σ*) for MPPCA and the proposed method, with a spatially varying noise level alone (left) and a spatial correlation alone (right). In each heatmap, columns sweep the spatial spread of the noise level std(*d*) and the correlation of a voxel with its nearest neighbor *ρ*(1), rows the number of volumes *N* and with it the matrix aspect ratio *N/M*, where *M* is the number of voxels in a patch. Typically, a dataset comes with *N* volumes and the patch is chosen to match, where the widely used patch size rule gives *M N* (Tournier et al., 2019). Instead, we fix the patch at 5×5×5 voxels, so *M* = 125 throughout and the aspect ratio varies through *N* alone. This keeps the imposed noise covariance the same across the grid, so the rows show how one covariance acts at different aspect ratios. Values are medians over voxels and noise draws. **(b)** Mean squared error (MSE) of the denoised series against the ground truth, as a percentage of the undenoised series’ MSE, over the same grid. All data was generated at thermal SNR = 20. The proposed method recovers the ground truth noise level across the grid and keeps the MSE low, whereas MPPCA falls short under correlation, increasingly so as the aspect ratio *N/M* grows.

**Figure S4:**
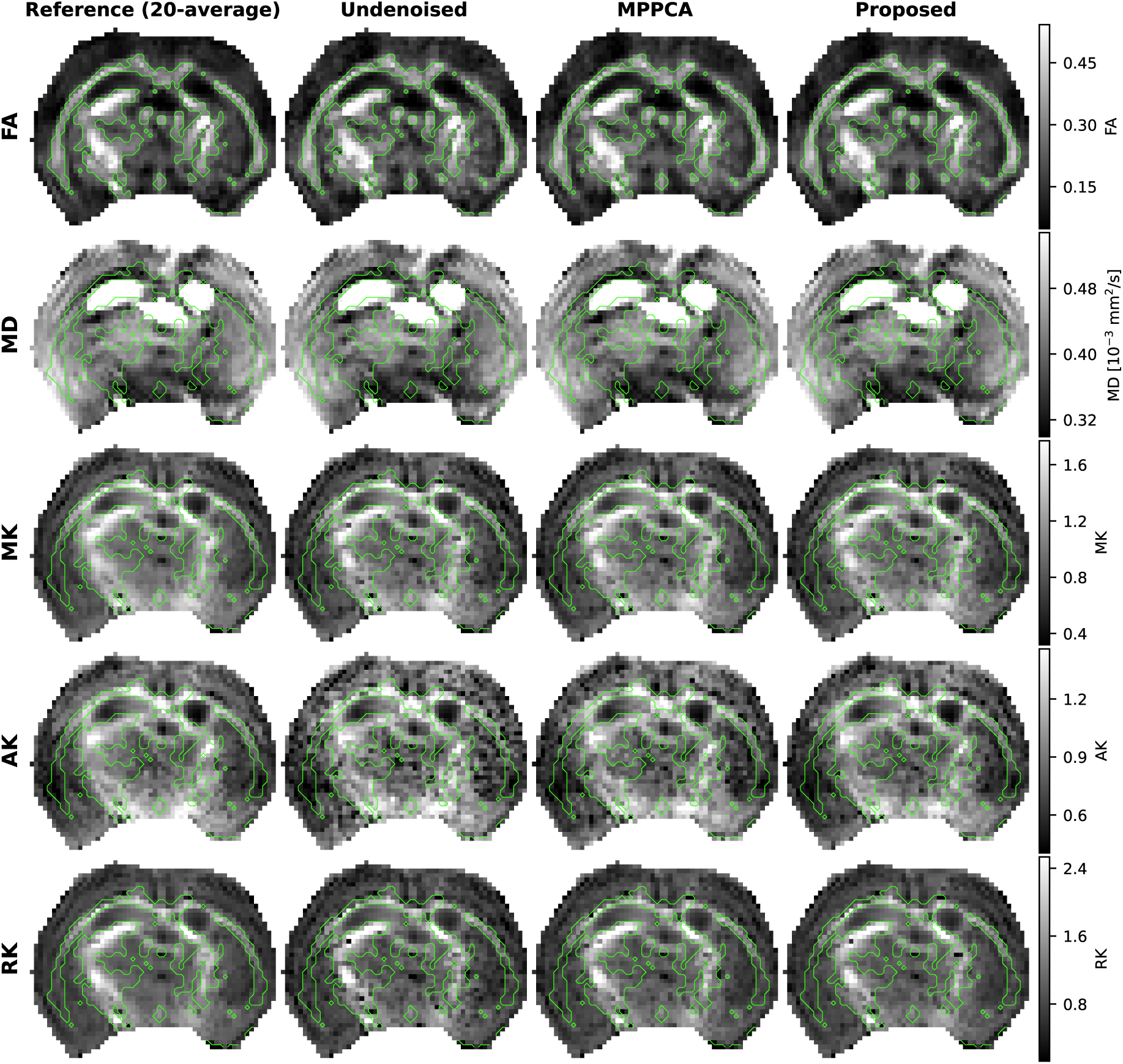
Ex vivo mouse brain dMRI: diffusion maps. Comparison of DTI and DKI maps for the slice shown in Figure 3a, with one row per metric and one column per series. The reference is fitted on the undenoised twenty-average volumes, whereas the remaining columns come from the two-average volumes, so the reference carries much higher SNR. DTI metrics include fractional anisotropy (FA) and mean diffusivity (MD); DKI metrics include mean, axial and radial kurtosis (MK, AK and RK). The green contours outline the white matter. Differences between the two denoising methods appear modest, while both reduce the speckle of the undenoised maps.

**Figure S5:**
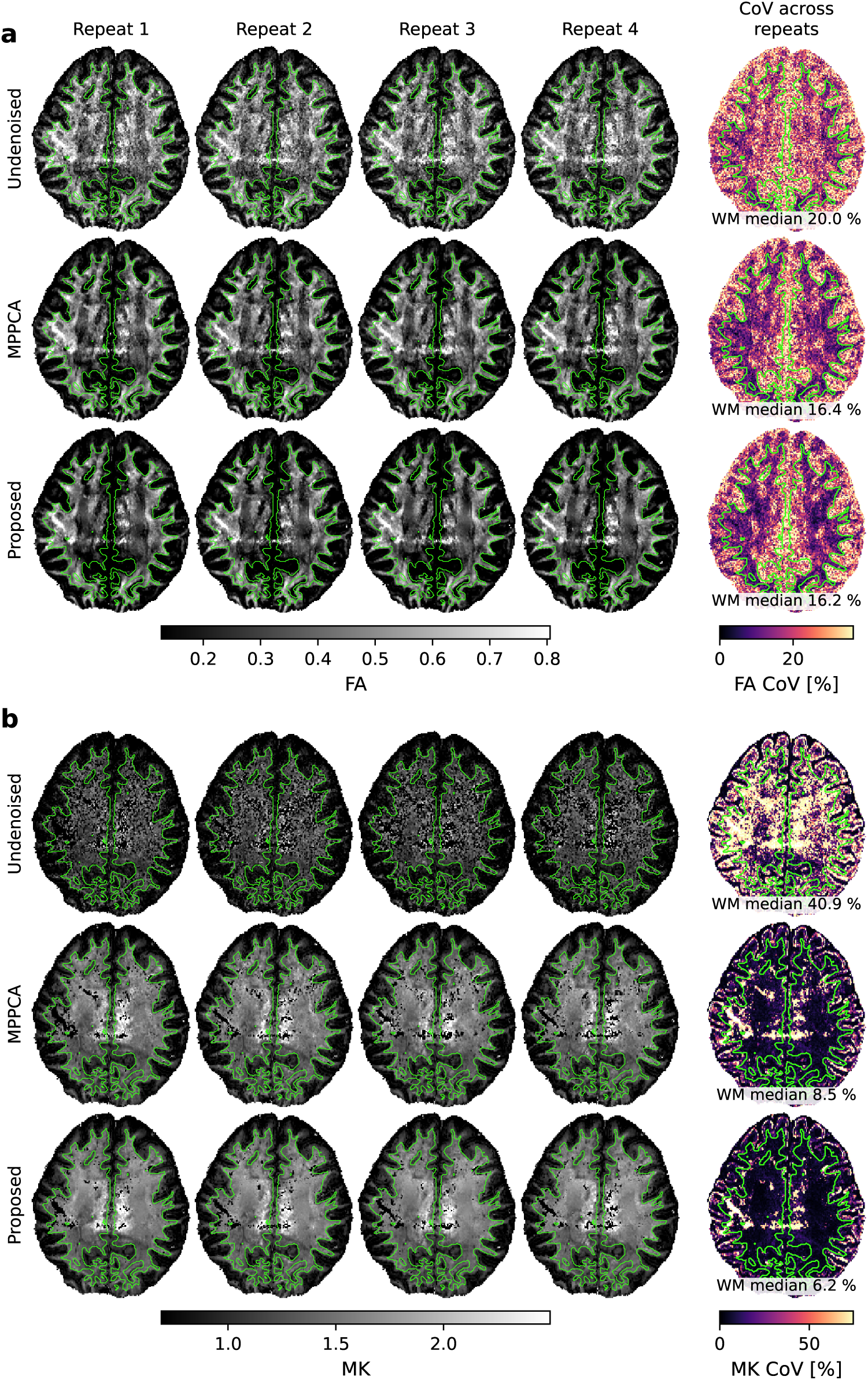
In vivo repeatability within one scanner at 0.9 mm: diffusion maps. Diffusion maps of **(a)** the DTI metric fractional anisotropy (FA) and **(b)** the DKI metric mean kurtosis (MK). The maps were computed from magnitude dMRI, corresponding to Figure 4a. Through (a–b), one column holds one repeat and one row one series. The last column gives each series’ coefficient of variation (CoV) across repeats per voxel, labeled with its median over the whole white matter. The green contours outline the white matter. The proposed method lowers the CoV in MK over MPPCA, while the FA CoV is similar.

**Figure S6:**
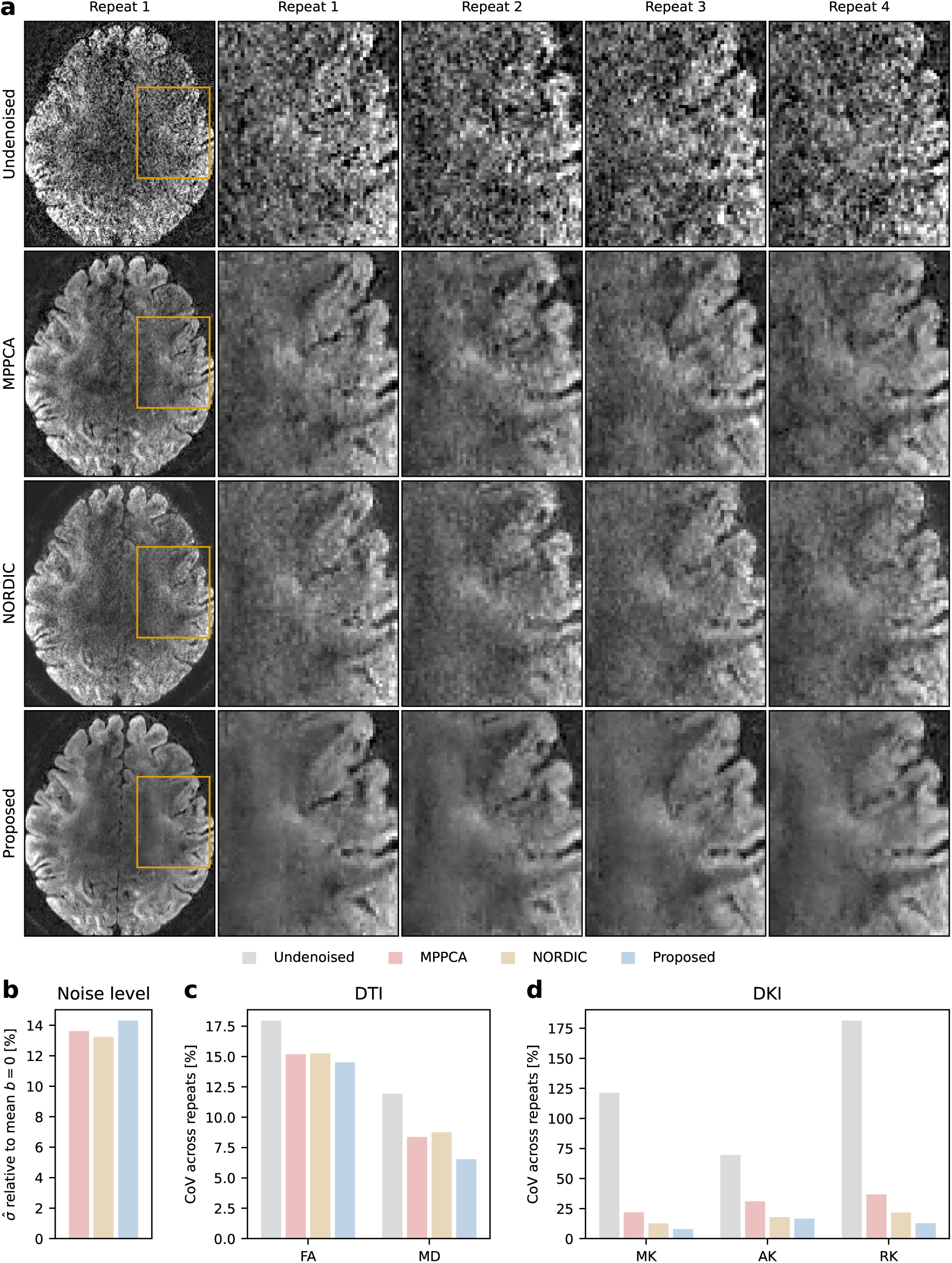
In vivo repeatability within one scanner at 0.9 mm: comparison with NORDIC. The figure repeats Figure 4 on complex data. All three denoising methods remove the background phase with the same step (Moeller, 2026; Sprenger et al., 2017). MPPCA and the proposed method then denoise the real part after rotation onto the real axis (Eichner et al., 2015), while NORDIC (Moeller et al., 2021) denoises the complex data directly. **(a)** Axial slice at *b* = 2000 s*/*mm^2^ across repeats, magnified on the region highlighted in the leading column. **(b)** Noise level *σ*^ estimated by each denoising method. **(c–d)** CoV for DTI and DKI metrics across repeats. Through (b–d), data are pooled over all voxels of the brain (b) or white matter (c–d). Bars give the median in (b). NORDIC estimated its noise map from MPPCA with 14×14×1 patches and denoised with 13×13×13, against 7×7×7 for MPPCA and the proposed method. The proposed method leaves less residual texture than MPPCA and lowers the CoV of MD and the DKI metrics, with NORDIC mostly in between. The FA CoV is similar across the denoising methods.

**Figure S7:**
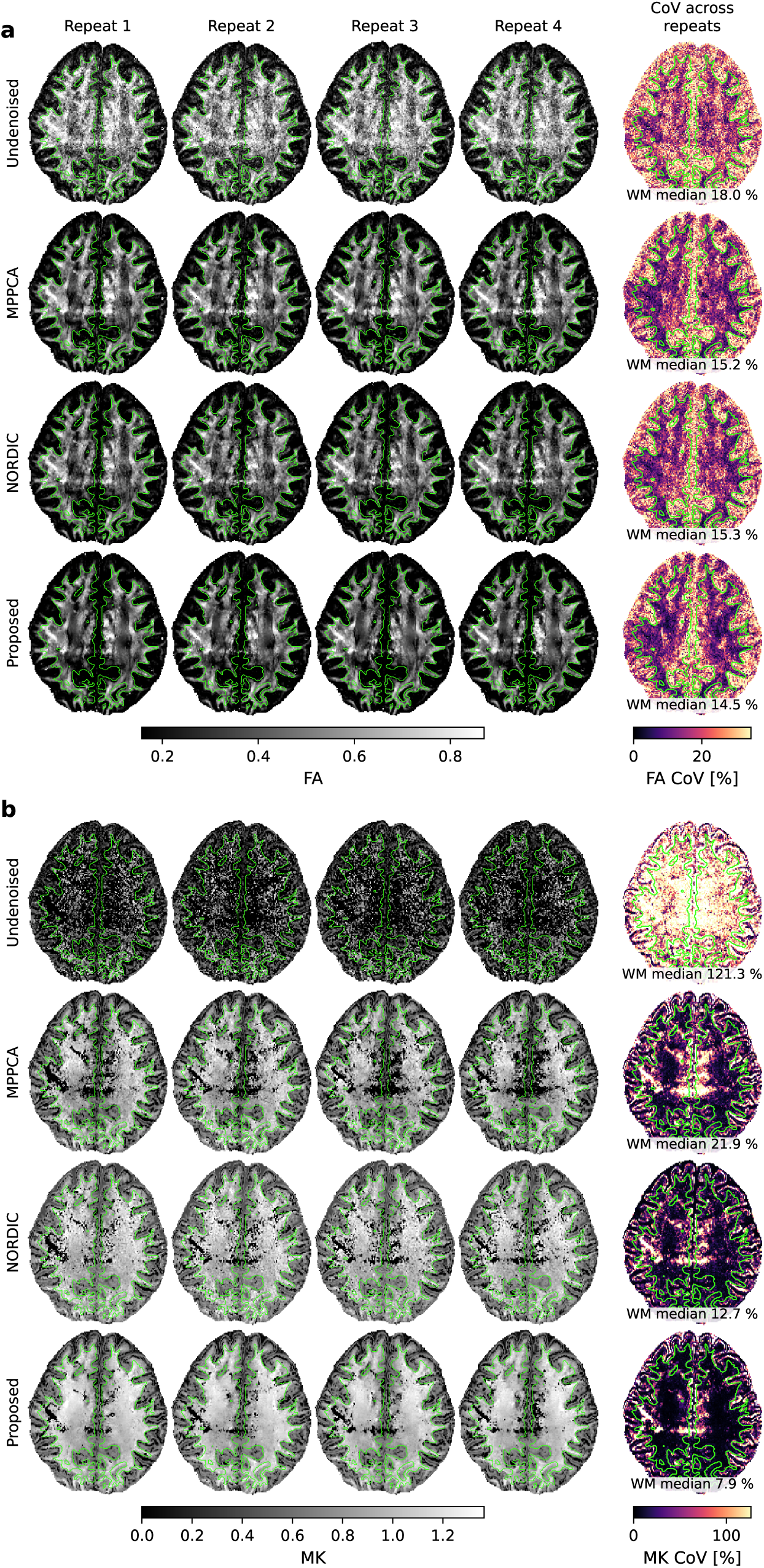
In vivo repeatability within one scanner at 0.9 mm: diffusion maps for the com-parison with NORDIC. Diffusion maps of **(a)** the DTI metric fractional anisotropy (FA) and **(b)** the DKI metric mean kurtosis (MK). The maps were computed from complex data, corresponding to Figure S6a. Through (a–b), one column holds one repeat and one row one series. The last column gives each series’ coefficient of variation (CoV) across repeats per voxel, labeled with its median over the whole white matter. The green contours outline the white matter. In MK, the proposed method gives the lowest CoV and NORDIC sits between it and MPPCA. In FA, the CoV is similar across the

**Figure S8:**
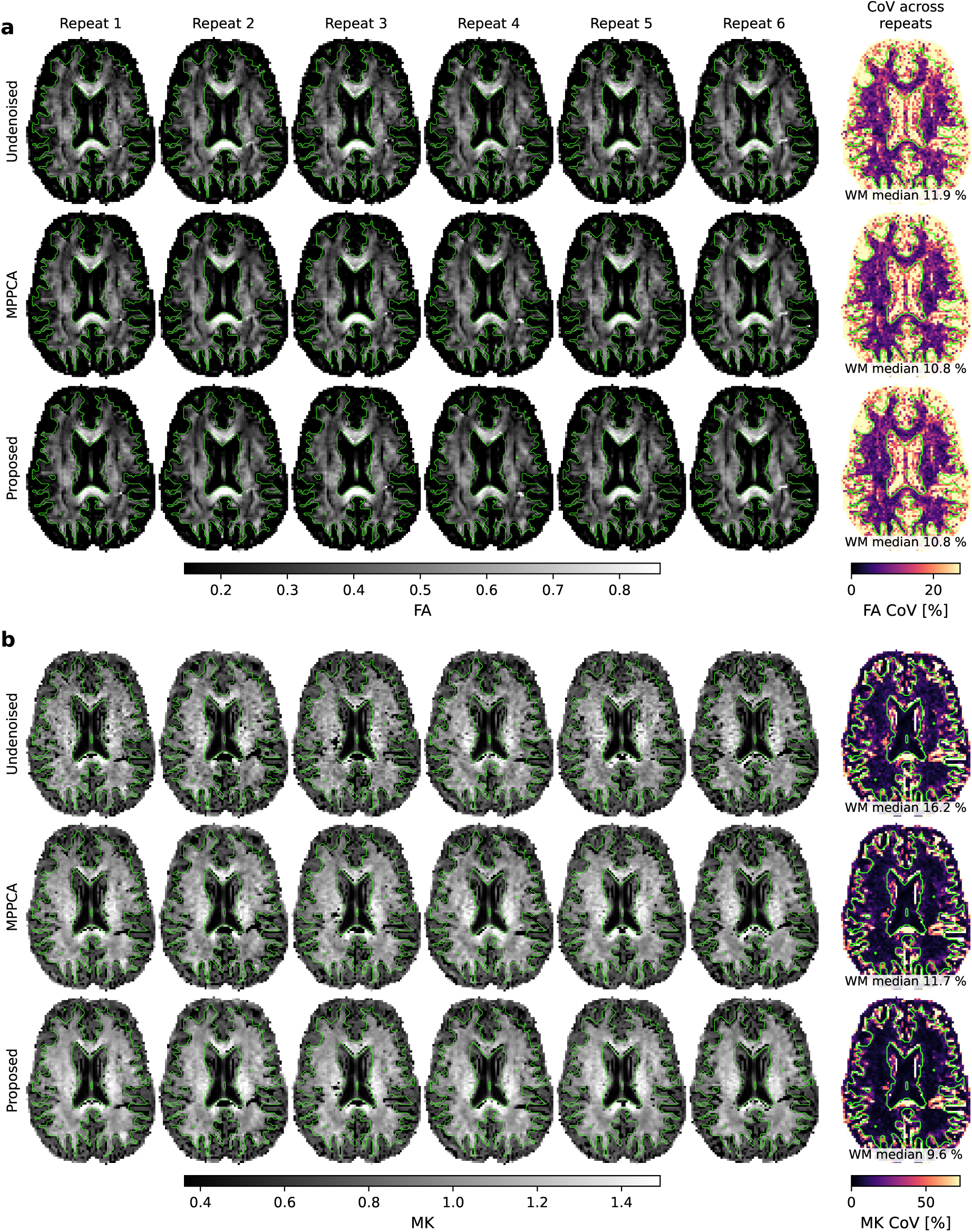
In vivo repeatability within scanners: diffusion maps. Diffusion maps of **(a)** the DTI metric fractional anisotropy (FA) and **(b)** the DKI metric mean kurtosis (MK). The maps were computed from magnitude dMRI, corresponding to Figure 5a. Through (a–b), one column holds one repeat and one row one series. The last column gives each series’ coefficient of variation (CoV) across repeats per voxel, labeled with its median over the whole white matter. The green contours outline the white matter. The proposed method lowers the CoV in MK over MPPCA, while the FA CoV is similar.

**Figure S9:**
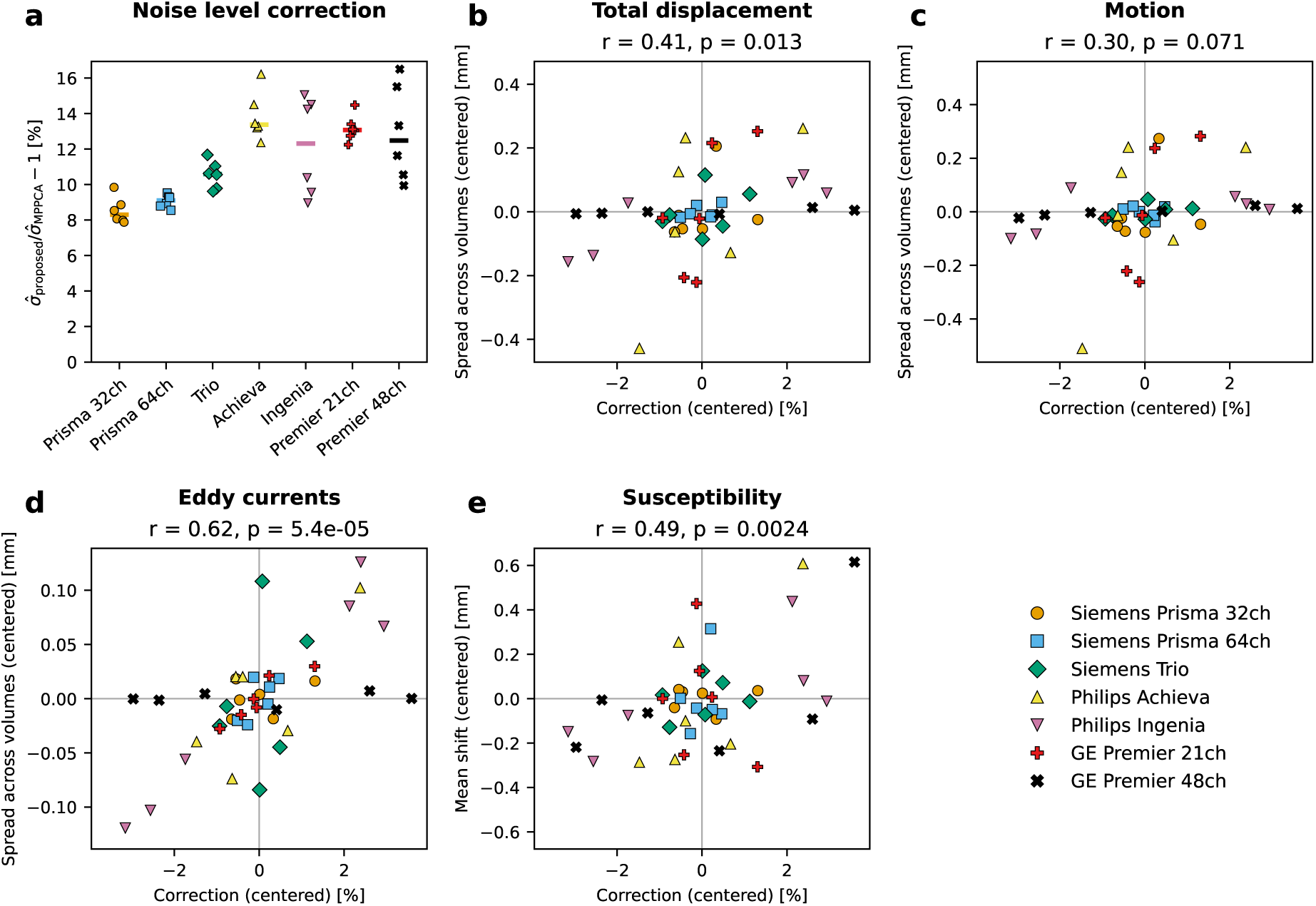
Association of the noise level correction with geometric displacement. **(a)** Correction of the proposed method relative to MPPCA, as the median over the brain voxels of a repeat. Each point denotes one repeat and the bar the median over the six repeats of a scanner. Subjects are distinct across scanners, so subject and scanner effects cannot be cleanly separated here. **(b–e)** Relationship between the correction and geometric displacement effects: total displacement (b), motion (c), eddy currents (d) and susceptibility (e). Panels (b–d) give the spread across the volumes of a repeat, whereas (e) gives the static shift along the phase encode direction, both estimated with FSL’s topup and eddy (Andersson & Sotiropoulos, 2016; Andersson et al., 2003; Jenkinson et al., 2012). All axes in (b–e) are centered on the scanner mean across its repeats. Above each panel, we report the repeated-measures correlation *r* (Bakdash & Marusich, 2017), which measures the association within scanners after accounting for across-scanner differences. The noise level correction appears most linked to the eddy current component, followed by susceptibility. However, the correction on the GE Premier 48ch varies without a matching change in any of the displacement measures.

**Figure S10:**
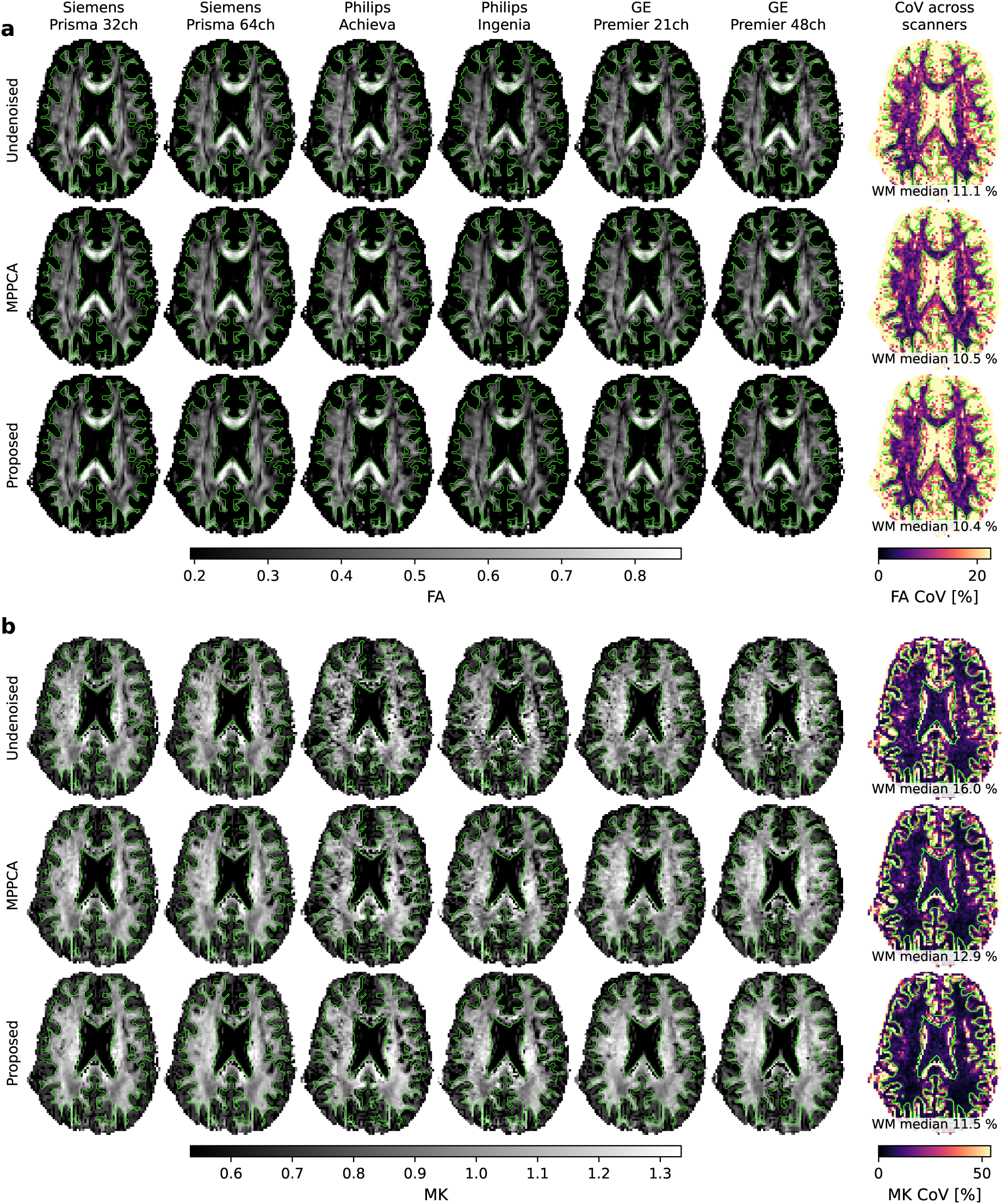
In vivo reproducibility across scanners: diffusion maps. Diffusion maps of **(a)** the DTI metric fractional anisotropy (FA) and **(b)** the DKI metric mean kurtosis (MK). The maps were computed from magnitude dMRI, corresponding to Figure 6a. Through (a–b), one column holds one scanner and one row one series. The last column gives each series’ coefficient of variation (CoV) across scanners per voxel, labeled with its median over the whole white matter. The green contours outline the white matter. The proposed method lowers the CoV in MK over MPPCA, while the FA CoV is similar.

**Figure S11:**
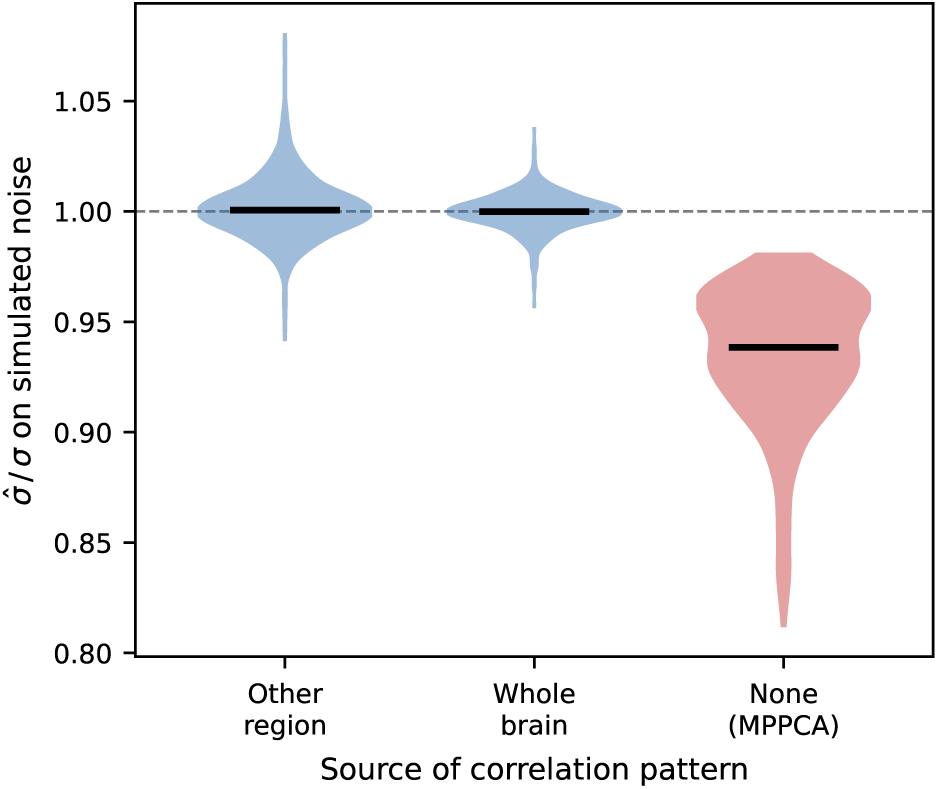
Correlation transfer between brain regions. Recovery of a known noise level on synthesized noise, after whitening with a correlation from another brain region of the same scan, or with the scan’s whole-brain correlation. Violins span the values with the median marked, and the dashed line sits at the known noise level. The last violin leaves the noise unwhitened for reference, as in MPPCA. The noise is synthesized with one region’s correlation, for each of 56 scans from 28 public datasets with two subjects each (see Section 2.6). Regions are the thirds of the brain mask along the phase encode, the readout and the slice axis, and each correlation is estimated by the proposed method on its region alone. Both the regional and the whole-brain correlation center on the known noise level, whereas MPPCA’s i.i.d. noise assumption underestimates it.

## Notes

### Competing Interest Statement

The authors have declared no competing interest.

